# Neuronal “parts list” and wiring diagram for a visual system

**DOI:** 10.1101/2023.10.12.562119

**Authors:** Arie Matsliah, Szi-chieh Yu, Krzysztof Kruk, Doug Bland, Austin Burke, Jay Gager, James Hebditch, Ben Silverman, Kyle Willie, Ryan Willie, Marissa Sorek, Amy R. Sterling, Emil Kind, Dustin Garner, Gizem Sancer, Mathias F. Wernet, Sung Soo Kim, Mala Murthy, H. Sebastian Seung, the FlyWire Consortium

## Abstract

A catalog of neuronal cell types has often been called a “parts list” of the brain, and regarded as a prerequisite for understanding brain function. In the optic lobe of *Drosophila*, rules of connectivity between cell types have already proven essential for understanding fly vision. Here we analyze the fly connectome to complete the list of cell types intrinsic to the optic lobe, as well as the rules governing their connectivity. We more than double the list of known types. Most new cell types contain between 10 and 100 cells, and integrate information over medium distances in the visual field. Some existing type families (Tm, Li, and LPi) at least double in number of types. We introduce a new Sm interneuron family, which contains more types than any other, and three new families of cross-neuropil types. Self-consistency of cell types is demonstrated through automatic assignment of cells to types by distance in high-dimensional feature space, and further validation is provided by algorithms that select small subsets of discriminative features. Cell types with similar connectivity patterns divide into clusters that are interpretable in terms of motion, object, and color vision. Our work showcases the advantages of connectomic cell typing: complete and unbiased sampling, a rich array of features based on connectivity, and reduction of the connectome to a drastically simpler wiring diagram of cell types, with immediate relevance for brain function and development.

## Introduction

Some of the greatest scientific discoveries of the 20th century concern the neural basis of sensory perception. Hubel and Wiesel’s discovery of simple and complex cells in the visual cortex not only entered neuroscience textbooks. The hypothetical neuronal wiring diagrams in their 1962 paper (Hubel and Wiesel 1962) inspired convolutional nets (Fukushima 1980; Y. LeCun et al. 1989), which eventually ignited the deep learning revolution in artificial intelligence (Yann LeCun, Bengio, and Hinton 2015). It may come as a surprise that no one has ever directly mapped wiring diagrams, however influential they may be. The best existing evidence requires the indirect inference of connectivity from cross-correlations in neural activity (Reid and Alonso 1995; Alonso and Martinez 1998), or the assumption that adjacency implies connectivity (Chapman, Zahs, and Stryker 1991).

Mapping the connections between individual neurons is still highly challenging in mammalian brains. Progress is being made by coupling calcium imaging *in vivo* with synaptic physiology in brain slices (Cossell et al. 2015) and calcium imaging of dendritic spines *in vivo* (Jia et al. 2010; Wilson et al. 2016). The reconstruction of a column of visual cortex from electron microscopic images is also becoming feasible (MICrONS Consortium et al. 2021; Schneider-Mizell et al. 2023). These are tiny slivers of visual systems; scaling up these approaches to tackle the full complexity of mammalian vision is still aspirational.

To imagine the future of mammalian visual neuroscience, it is helpful to extrapolate from a brain of more modest size, that of the fly, for which a complete neuronal wiring diagram is now just a few keystrokes away. Especially over the past 15 years, visual neural circuits have been intensively investigated in *Drosophila* (Currier, Pang, and Clandinin 2023) with great progress in understanding the perception of motion (Shinomiya et al. 2022; Borst and Groschner 2023), color (Schnaitmann, Pagni, and Reiff 2020), and objects (Wu et al. 2016), as well as the role of vision in complex behaviors like courtship (Ribeiro et al. 2018). The recent release of a neuronal wiring diagram of a *Drosophila* brain (Zheng et al. 2018; Dorkenwald, Matsliah, et al. 2023; Schlegel et al. 2023) poses an unprecedented opportunity. The first wiring diagram for a whole brain contains as a corollary the first wiring diagram for an entire visual system, as well as all the wiring connecting the visual system with the rest of the brain.

Most visual neurons, and more generally most neurons, are situated in the optic lobes of the *Drosophila* brain (Fig. S1a). In the reconstructed brain (Dorkenwald, Matsliah, et al. 2023), about 38,500 neurons are intrinsic to the right optic lobe, meaning that their synapses are fully contained in this region. A wiring diagram for the *Drosophila* visual system must include this many neurons at least. While minuscule compared to a mammalian visual system, this is still too complex to comprehend or even visualize. It is essential to reduce the complexity by describing the connectivity between types of cells. For example, the roughly 800 ommatidia in the compound eye send photoreceptor axons to roughly 800 L1 cells in the lamina, which in turn connect with roughly 800 Mi1 cells. That is a lot of cells and connections, but they can all be described by the simple rules that photoreceptors connect to L1, and L1 connects to Mi1. Some of these connectivity rules have long been known (Meinertzhagen and O’Neil 1991; Gao et al. 2008), and more have been discovered over the past decade (S.-Y. Takemura et al. 2013, 2015; S. Takemura et al. 2017; Shinomiya et al. 2019, 2022), but this knowledge is fragmentary and incomplete.

In the present work, we exhaustively enumerate all cell types intrinsic to the optic lobe, and find all rules of connection between them. We have effectively collapsed 38,500 intrinsic neurons onto just 226 types, a more than 150× reduction. The wiring diagram is reduced from a 38,500×38,500 matrix to a 226×226 matrix, an even huger compression. We additionally provide rules of connectivity between intrinsic types and types of boundary neurons, defined as those that connect the optic lobe with regions in the central brain. Such central brain regions are generally multimodal and/or sensorimotor, mixing information coming from the eyes and other sense organs, so we regard the optic lobe proper as the fly’s visual system.

In our connectomic approach, a cell type is defined as a set of cells with similar patterns of connectivity (Scheffer et al. 2020), and this definition turns out to be critical for distinguishing between cell types that are morphologically very similar. Because connectivity is so closely related to function, cells of the same connectomic type are expected to share the same function (H. Sebastian Seung and Sümbül 2014). By the same logic, cell types with similar patterns of connectivity ought to have similar functions. We support this claim by dividing our cell types into clusters, and interpreting the functions of these clusters in terms of motion, object, and color vision. This provides preliminary clues concerning the functions of newly discovered cell types, as well as the previously known cell types for which functional information has been lacking.

The present work is based on version 783 of the FlyWire connectome, which incorporates proofreading up to September 30, 2023. Our work will continue to be updated as new versions of the connectome are released (see Discussion). All information can be browsed, searched, and downloaded at the FlyWire Codex (codex.flywire.ai). This addition to the existing array of FlyWire resources (Dorkenwald, Matsliah, et al. 2023; Schlegel et al. 2023) will accelerate the pace of discovery for many researchers.

### Class, Family, and Type

Neurons intrinsic to the optic lobe are those with synapses entirely contained inside the optic lobe, and are the main topic of this study (Fig. S1a). In addition, there are “boundary” neurons that straddle the optic lobe and the rest of the brain (Fig. S1b). Boundary neurons fall into several classes. Visual projection neurons (VPNs) have dendrites in the optic lobe and extend axons to the central brain. Visual centrifugal neurons (VCNs) do the opposite. Heterolateral neurons are intrinsic to the pair of optic lobes; they extend from one optic lobe to the other while making few or no synapses in the central brain. Boundary neurons have been divided into types by (Schlegel et al. 2023).

Our work is based on the brain of a single *Drosophila* adult female, which was reconstructed by the FlyWire Consortium (Dorkenwald, Matsliah, et al. 2023). We have proofread roughly 38,500 intrinsic neurons in the right optic lobe, as well as 4,000 VPNs, 250 VCNs, 200 heterolateral neurons, and 4700 photoreceptor cells. 80% of the synapses of intrinsic neurons are with other intrinsic neurons, and 20% are with boundary neurons. These statistics suggest that the optic lobe communicates more with itself than with the rest of the brain.

We divide optic lobe intrinsic neurons into four broad classes. Cells of the columnar class (Fig. 1a) have axons oriented parallel to the main axis of the visual columns (see Methods for definition of “axon”). Following (Fischbach and Dittrich 1989), the arbor of a columnar neuron is allowed to be wider than a single column; what matters is the orientation of the axon, not the aspect ratio of the arbor. Photoreceptor or retinula cells are columnar but are not intrinsic to the optic lobe, strictly speaking, because they enter from the retina. Nevertheless, they will sometimes be included with intrinsic types in the analyses and comparisons to follow.

**Figure 1.**
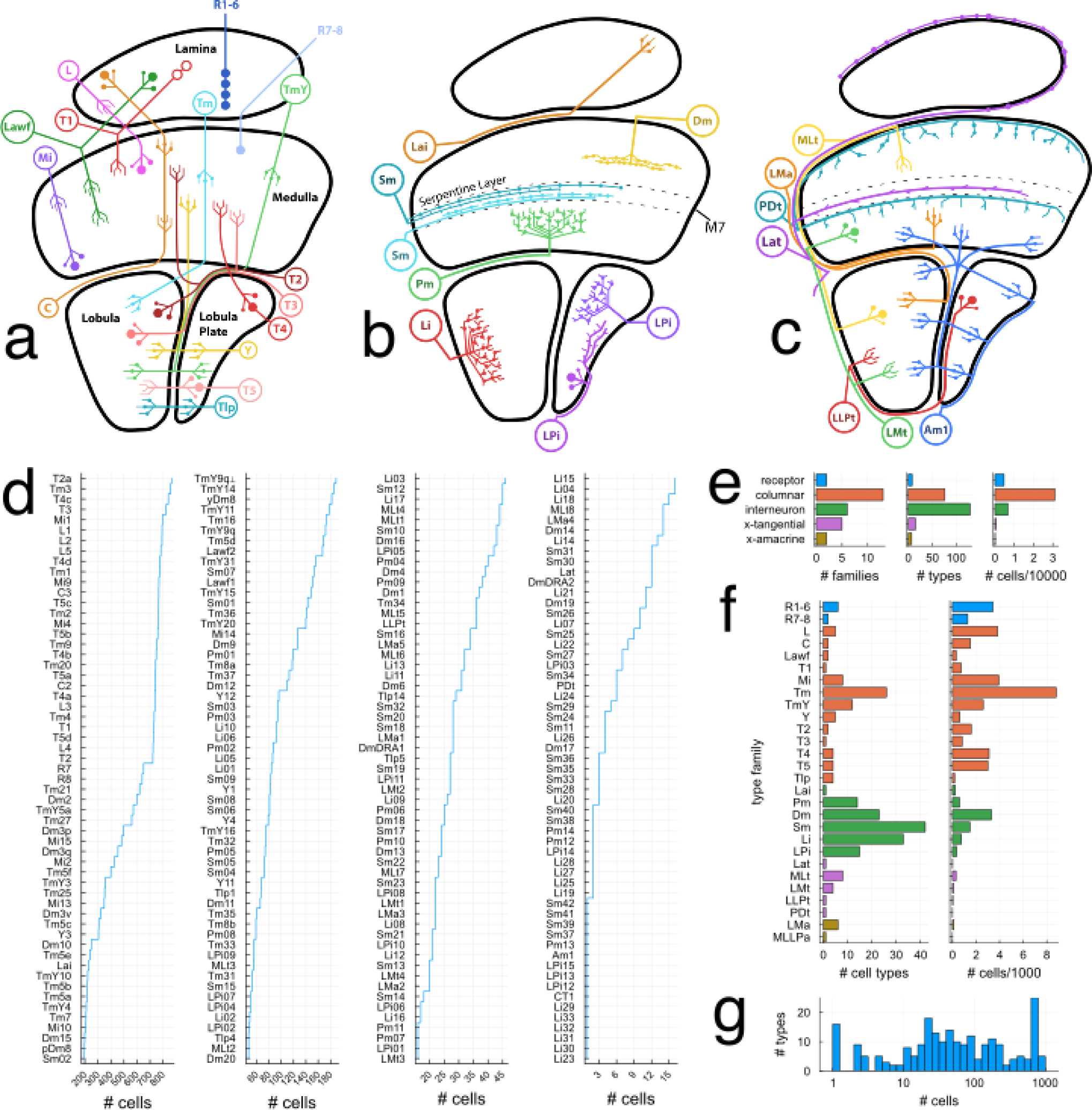
Class, family, type, and cell. (a) Families in the columnar class. (R: Retinula, L: Lamina monopolar, C: Centrifugal, Lawf: Lamina wide-field, Mi: Medulla intrinsic, Tm: Transmedullary, TmY: Transmedullary Y, Tlp: Translobula plate) (b) Families in the interneuron class. Sm is novel. (Lai: Lamina intrinsic, Pm: Proximal medulla, Dm: Distal medulla, Sm: Serpentine medulla, Li: Lobula intrinsic, LPi: Lobula Plate intrinsic). (c) Families in the cross-neuropil tangential and amacrine classes. For tangential families, axon and dendrite are distinguished graphically. All are novel except Lat. PDt and MLLPa are not shown for clarity. (MLt: Medulla Lobula tangential, LLPt: Lobula Lobula Plate tangential, LMt: Lobula Medulla tangential, LMa: Lobula Medulla amacrine, Lat: Lamina tangential, PDt: Proximal to Distal medulla tangential. A: Anterior. L: Lateral. V: Ventral) (d) Cell types ordered by number of cells in each type, starting with the most numerous types. Cell counts are based on v783. (e) Number of families (left), types (middle), and cells (right) in each class. (f) Number of types (left) and cells (right) in each neuropil-defined family. Bold font indicates families that are entirely new, or almost entirely new. MLLPa (Medulla Lobula Lobula Plate amacrine) is a synonym for Am1. (g) Number of types versus number of cells in a type. X-axis denotes type size (log-scale), and Y-axis the number of types with matching size. The peak near 800 consists of the “numerous” types, those with approximately the same cardinality as the ommatidia of the compound eye.

The optic lobe (Fig. S1a, b) contains four main neuropils (lamina, medulla, lobula, and lobula plate) and a smaller fifth neuropil (accessory medulla). Following (Fischbach and Dittrich 1989) we further distinguish between distal and proximal medulla, regarding them as two separate neuropils (Fig. S1c). A columnar cell spans multiple neuropils (Fig. 1a). Cells of the local interneuron class (Fig. 1b) are defined as being confined to a single neuropil. We also define two classes that cross multiple neuropils but are not columnar. A cross-neuropil tangential cell (Fig. 1c) has an axon that is oriented perpendicular to the main axis of the visual columns as it runs inside a neuropil. A cross-neuropil amacrine cell (Fig. 1c) lacks an axon. Interneurons are typically amacrine, but sometimes have an axon in the tangential orientation.

Each class is divided into multiple families. A family is defined as a set of cells that share the same neuropils (Fig. 1a, b, c, Methods). For example, the Tm family projects from the distal medulla to the lobula, while the TmY family projects from the distal medulla to both the lobula and lobula plate (Fig. 1a). (Tm and TmY also typically receive inputs in the proximal medulla.)

Each family is divided into cell types. All 226 intrinsic types as well as photoreceptor types are available for 3D interactive viewing at the FlyWire Codex (codex.flywire.ai). Table S1 is a master list of all types and their properties. Data S1 contains one “card” for each type, which includes its discriminative logical predicate (see below), basic statistics, diagram in the style of (Fischbach and Dittrich 1989) showing stratification and other single-cell anatomy, and 3D renderings of all the cells in the type.

Most neurons in the optic lobe are columnar (Fig. 1e, right), and half of the families are columnar (Fig. 1e, left). Interneurons constitute just 17% of optic lobe intrinsic neurons, but the majority of cell types (Fig. 1e, middle). A columnar family (Tm) contains more cells than any other family (Fig. 1f, right). An interneuron family (Sm) contains more types than any other family (Fig. 1f, left).

The columnar families (Fig. 1a) were defined by (Fischbach and Dittrich 1989). The Sm interneuron family is new (Fig. 1b), and some of the cross-neuropil families are wholly or almost wholly new (Fig. 1c). Over half of the cell types are new, and many of these are interneuron types.

### Connectomic approach to cell types

For each cell, we define an output feature vector by the number of output synapses onto neurons of cell type *t*, which runs from 1 to *T*. This is a *T*-dimensional vector, where *T* is the number of cell types. The output feature vector is a row of the cell-to-type connectivity matrix (Methods). For each cell, we similarly defined an input feature vector by the number of input synapses received from neurons of cell type *t*. This is a column of the type-to-cell connectivity matrix (Methods). Alternatively, feature vectors can be based on connection number, where a connection is defined as two or more synapses from one neuron to another (Methods), and this gives similar results.

The input and output feature vectors were concatenated to form a 2*T*-dimensional feature vector. If we include only intrinsic types (Fig. 2a), then *T* is 226 and the dimensionality 2*T* is 452. If we include both intrinsic and boundary types, then *T* is over 700 and the dimensionality 2*T* is over 1400 (not shown). We experimented with both dimensionalities, and as will be explained below, the results do not depend on the boundary types. The lower dimensionality feature vector with only intrinsic types is sufficient for defining types.

**Figure 2.**
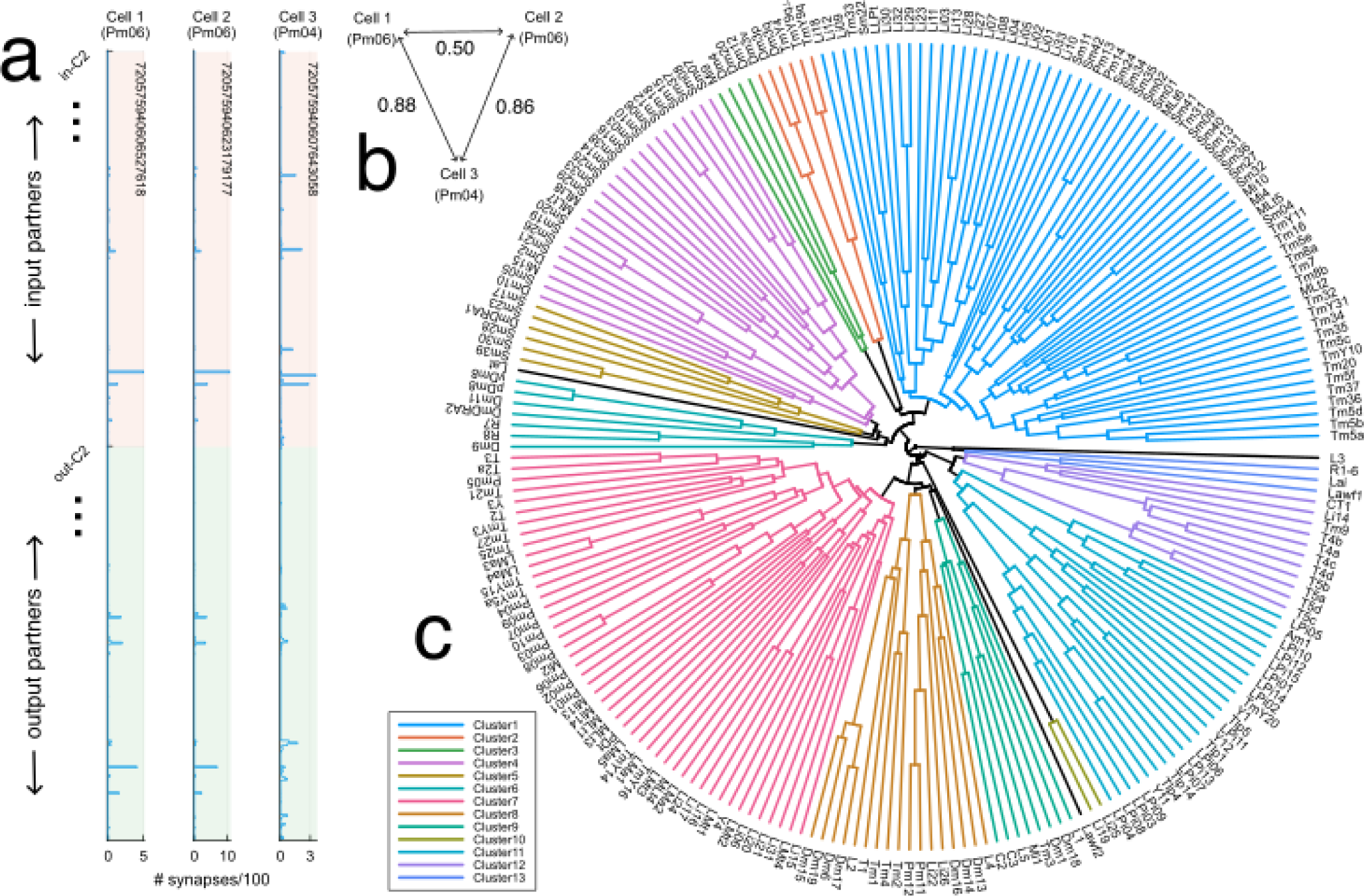
Clustering of cells and cell types based on connectivity. (a) Feature vectors for three example cells. The horizontal axis indicates the synapse numbers the cell receives from presynaptic types (red region of vertical axis) and sends to postsynaptic types (green region of vertical axis). Cells 1 and 2 (same type) have more similar feature vectors to each other than to Cell 3 (different type). (b) Cells 1 and 2 (same type) are closer to each other than to Cell 3 (a different type), according to weighted Jaccard distances between the cells’ feature vectors. Such distances are the main basis for dividing cells into cell types (Methods). (c) Dendrogram of cell types. Cell types that merge closer to the circumference are more similar to each other. Flat clustering (13 colors) is created by thresholding at 0.91. A few clusters containing single types are uncolored. To obtain the dendrogram, feature vectors of cells in each type were summed or averaged to yield a feature vector for that cell type, and then cell type feature vectors were hierarchically clustered using average linkage. Jaccard distances run from 0.4 (circumference) to 1 (center).

A cell type is defined as a set of cells with similar feature vectors (Scheffer et al. 2020). Cells of the same type are near each other in feature space, while cells of different types are far away (Fig. 2b). This was quantified using weighted Jaccard distance, which we will call Jaccard distance for short (Methods).

In order to define feature vectors, some cell types must already exist. An initial set of cell types was defined by human analysts employing traditional morphological criteria (Methods). Namely, each neuropil was divided into multiple layers (Fischbach and Dittrich 1989) (Fig. S1c). A cell type was classically defined as a set of neurons that stratify across the layers in the same way. Such cell types were used to compute feature vectors, and then hierarchical clustering was applied. In many cases, this led to further division into cell types that could not be distinguished by traditional criteria. In other cases, it led to grouping of morphological variants into a single type. After splitting or merging types, the feature vectors were recomputed and the process continued iteratively.

The final cell types are validated in several ways (Methods). We show that our clustering is self-consistent, in the sense that almost all cells end up in the original cluster if we attempt to reassign each cell’s feature vector to the nearest cluster. For more interpretable evaluations, we construct compact connectivity-based discriminators that can predict cell type membership (Fig. S2, Data S2). We show that membership can be accurately predicted by a logical conjunction of on average five synaptic partner types. For each intra-neuropil type, we also provide a 2D projection of the connection matrix that can be used to discriminate that type from others in the same neuropil (Fig. S3c, Data S3).

### Hierarchical clustering of cell types

We defined a connectomic cell type as a set of cells with similar feature vectors based on connectivity. It follows that cells of the same type should share the same function, if one accepts the maxim, “Nothing defines the function of a neuron more faithfully than the nature of its inputs and outputs” (Mesulam 2005).

Taking one step further, the same maxim implies that cell types with similar feature vectors should have similar visual functions. A cell type feature vector can be obtained by summing the feature vectors over all cells in that type. We separately sum and normalize the input and output feature vectors, and then concatenate the results.

We computed the Jaccard distance between all pairs of cell type feature vectors. It is hard to extract insights directly from such a large distance matrix, so we used it to compute a hierarchical clustering (Fig. 2c). The dendrogram was cut to yield a flat clustering, which is indicated by coloring the dendrogram (Fig. 2c). Each cluster contains some cell types for which visual functions are known. Later on, we will extrapolate from this information to assign hypothetical functions to entire clusters.

The precise memberships in the clusters warrant cautious interpretation, as the clusters are the outcome of just one clustering algorithm (average linkage), and differ somewhat if another clustering algorithm is employed. Each cluster contains core groups of types that are highly similar to each other, i.e. types that merge in the dendrogram farther from the origin. These are more certain to have similar visual functions, and tend to be grouped together by any clustering algorithm. Types that are merged closer to the origin are less similar, and their cluster membership is more arbitrary. Some degree of arbitrariness is inevitable when one divides the visual system into separate subsystems, because subsystems interact with each other, and types that mediate such interactions are borderline cases.

Each cluster is generally a mixture of types from multiple neuropil families. Skeptics might regard such mixing as arising from the “noisiness” in the clustering noted above. Indeed, the nearest types, those that merge in the dendrogram farther from the center (Fig. 2c), tend to be from the same neuropil family. But plenty of dendrogram merges between types of different families happen at intermediate distances rather than the largest distances. Therefore some of the mixing of types from different neuropil families seems genuinely rooted in biology.

### Type-to-type connectivity

We define a type-to-type connection matrix in which the *st* element is the number of synapses from cell type *s* to cell type *t* (Methods). The type-to-type matrix is drastically smaller than the cell-to-cell matrix. Nevertheless, it is still quite large, presenting challenges to visualization and understanding. Figure S4 attempts to show the entire matrix. The area of each dot encodes the number of synapses in some type-to-type connection. Dot area saturates above 3600 synapses, in order to make weaker connections visible. The numerical values of the matrix elements are provided in the Supplementary Information, and can be downloaded from the Codex.

The type-to-type connection matrix can also be visualized as a directed graph. Since showing all connections is visually overwhelming, it is important to find ways of displaying meaningful subsets of connections. One that we have found helpful is to display the top input and output connections of each type (Fig. 3, S5, S6). In such a graph, every node must have at least one outgoing and one incoming connection. But some nodes can still be “hubs” with many visible connections. For example, Mi1 is the top input to a large number of postsynaptic types.

**Figure 3.**
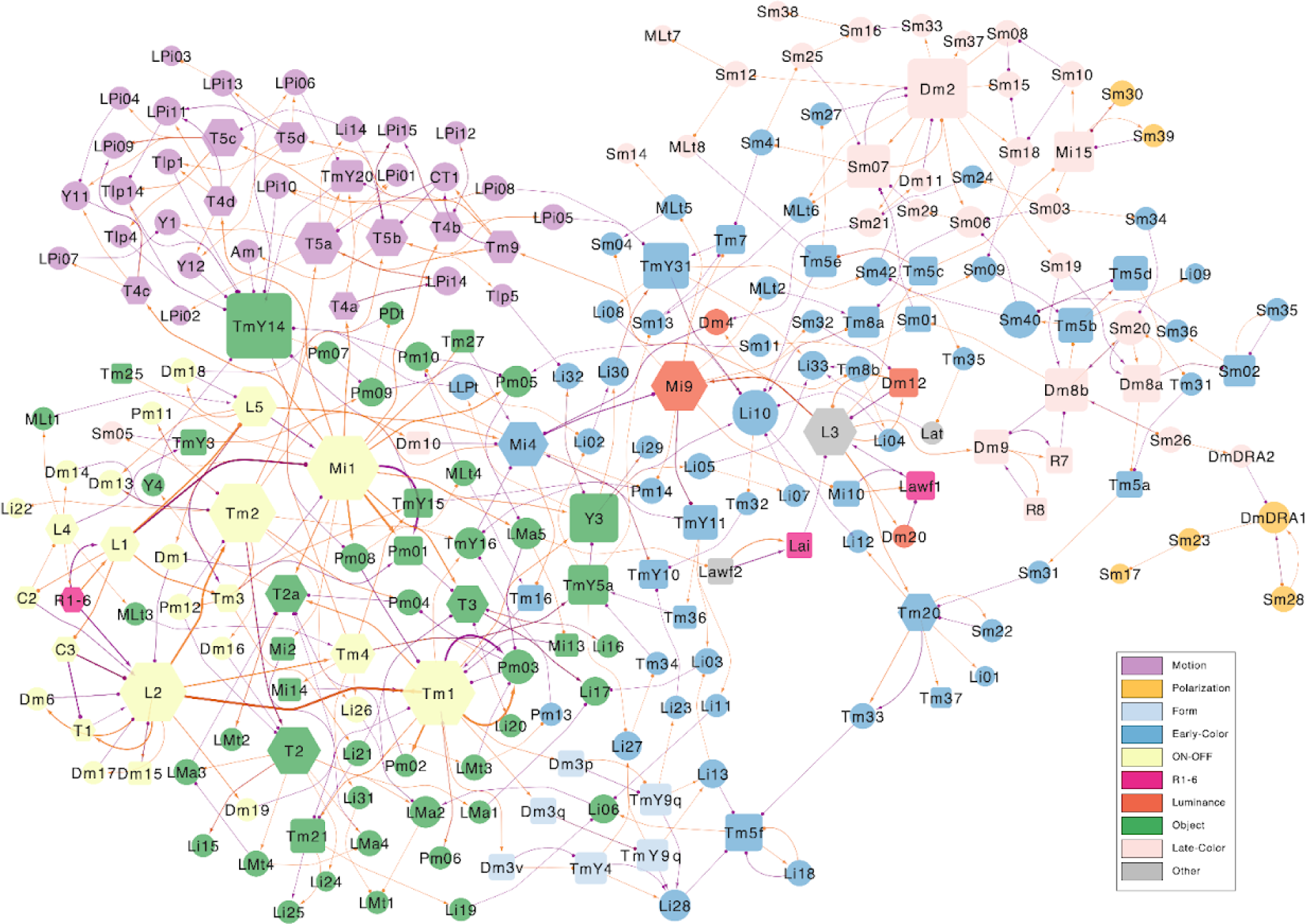
Wiring diagram of cell types (top input and output connections) A simplified wiring diagram of all cell types intrinsic to the optic lobe, as well as photoreceptors. For clarity, only the top input and output connections of each type are drawn. Node size encodes the number of drawn connections, so that “hub” types look larger. Node color indicates membership in the subsystems defined in the text (see legend). Node shape indicates number of cells (hexagons 800+, rectangles 100-799 and circles 1-99). Orange edges indicate “top input” relationships, purple edges indicate “top output”. Arrow tips indicate excitation and circle tips indicate inhibition.

The nodes of the graph were positioned in 2D space by a graph layout algorithm that tends to place visibly connected types close together. It turns out that nearby nodes in the 2D graph layout space tend to belong to the clusters that were extracted from the high dimensional connectivity-based feature vectors (compare node colorings of Fig. 3 with clusters of Fig. 2).

We can also normalize the type-to-type connection matrix to be the fraction of synapses from cell type *s* to cell type *t*. Depending on the normalization, this could be the fraction of input to type *t* or fraction of output from type *s* (Methods). Input and output fractions are shown in Data S4, and are equivalent to the cell type feature vectors defined earlier. Each column of the heatmap contains the input or output fractions of the corresponding cell type listed at the bottom of the heatmap. The rows include all partners that contribute at least 0.02 input or output fraction to any of the reference types listed at the bottom of the heatmap. The heatmaps include partner boundary types as well as intrinsic types. The colormap has been chosen to saturate at 0.2, in order to make small values visible, so large values (≥0.2) are indistinguishable. The viewer should be alert to artifacts that can arise due to normalization. If a cell type makes very few synapses overall, that can result in large input and output fractions, giving the mistaken impression of connections that are strong when they are actually weak or nonexistent.

### Perplexity as a measure of degree of connectivity

As mentioned above, some cell types are hubs with a high degree of connectivity to other types. The degree of a node can be defined as the number of nodes to which it is connected. The elements of the type-to-type connection matrix vary greatly in magnitude, and it is unclear whether weak connections should be included when computing degree. This can be handled by thresholding the connection matrix before computing degree. For a threshold-independent measure, however, we instead calculate a “perplexity” for each node. The outgoing connection strengths (synapse counts) are normalized as if they were a probability distribution, and out-perplexity is defined as the exponential of the entropy of this distribution. Out-perplexity reduces to out-degree in the special case that the distribution is uniform over the connected partners. In-perplexity is defined analogously.

If cell types are ranked by product of out- and in-perplexity (Fig. S7a), then TmY5a is the most connected hub, and various types in the lamina and distal medulla are the least hub-like. Motion-related cell types generally do not have high perplexity. Out-perplexity tends to be greater than in-perplexity (Fig. S7a), though they are positively correlated (Fig. S7b).

One might expect that “early” types in visual processing would have divergent connectivity, to distribute photoreceptor signals to many targets, while “late” types would have convergent connectivity, summarizing the final results of optic lobe computations for use by the central brain. This idea can be tested by ranking types according to ratio of out- to in-perplexity (Fig. S8). Indeed, the top of the list includes early types like the inner photoreceptors R7 and R8, L3 and L5, and many Dm and Pm interneuron types. And many Sm types are near the bottom of the list, as befits their extensive connectivity with VPNs in the serpentine layer. But reality is clearly more complex than the simple idea that motivated the graph.

### The “numerous” cell types

The most numerous types contain about 800 cells, the same as the estimated number of ommatidia (Fig. 1d). The least numerous types contain only a single cell. The number of cells per type is multimodally distributed (Fig. 1g), with two sharp modes at 1 and roughly 700. Most of the distribution lies between these modes (Fig. 1g, S1d).

The most numerous types have long been known (Fischbach and Dittrich 1989). It is the less numerous types where our knowledge has been incomplete, and arguably they are where much of the magic of vision happens. As with the photoreceptors, neural activity in the most numerous cell types like L1 and Mi1 mostly encodes information about the image at or near single points in visual space. But perception requires the integration of information from points that can be quite distant from each other, and this is done by the larger neurons that belong to the less numerous types.

Photoreceptor axons originating in the roughly 800 ommatidia of the compound eye project retinotopically to the lamina and medulla neuropils. The medulla is divided into columns, which are presumed to be in one-to-one correspondence with the ommatidia. (S.-Y. Takemura et al. 2013) defined “modular” cell types as those that are in one-to-one correspondence with columns. Based on their reconstruction of seven medulla columns, (S.-Y. Takemura et al. 2015) found that there were 20 modular types. These largely correspond to the cell types that contain from 720 to 800 cells in v783 (Fig. 1d).

The top end (800) of this range is likely the true number of columns in this optic lobe. In the lower end of this range, the deficit relative to 800 could be due to underrecovery of cells (Methods) so our data is still consistent with modularity. R7 and R8 are about 650 cells each, but this is not inconsistent with modularity because photoreceptors are especially challenging to proofread in this dataset (Methods). Tm21 (a.k.a. Tm6), Dm2, TmY5a, Tm27, and Mi15 have much lower numbers, so we agree with (S.-Y. Takemura et al. 2015) that they are not modular.

On the other hand, some of our types (T2a, Tm3, T4c, T3) contain more than 800 proofread cells (Fig. 1d), which violates the definition of modularity. This partially agrees with (S.-Y. Takemura et al. 2015), who regarded T3 and T2a as modular, and T4 and Tm3 as not modular. T4 is an unusual case, as T4c is above 800 while the other T4 types are below 800, though these numbers could still creep upward with further proofreading.

A genuine analysis of modularity requires going beyond simple cell counts, and analyzing locations in order to check the idea of one-to-one correspondence. Such an analysis is left for future work. For now, we will apply the term “numerous” to those types containing 720 or more cells, as well as R7 and R8, and do not commit to whether these types are truly modular.

(S.-Y. Takemura et al. 2015) provide a matrix of connections between their modular types. This shows good agreement with our data (Fig. S9, Methods), and validates our reconstruction in the optic lobe. This validation complements the estimates of reconstruction accuracy in the central brain that are provided in the flagship paper (Dorkenwald, Matsliah, et al. 2023).

The connectivity and visual responses of the numerous types are largely known from previous work. In the following, we will provide functional interpretations of the cell type clusters shown in the dendrogram of Fig. 2c. It turns out that the numerous types that belong to a single cluster have similar functions, which enables us to ascribe a function to each cluster as a whole. In other words, we extrapolate from the functions of the numerous types to yield preliminary clues regarding the functions of the less numerous types.

### ON, OFF, and luminance channels

We propose that Cluster8 and Cluster9 be regarded as OFF and ON channels, respectively, which are dedicated to carrying information about light decrements (“OFF” stimuli) and light increments (“ON” stimuli). Our concept is similar to the well-known ON and OFF motion pathways, but differs because our ON and OFF channels are general-purpose, feeding into the object and color subsystems as well as the motion subsystem.

Cluster8 contains the OFF cells L2, L4, Tm1, Tm2, and Tm4. Cluster9 contains the ON cells L1, L5, Mi1, and Tm3 and also the OFF cell L1. It makes sense to assign L1 to the ON channel even though it is an OFF cell, because L1 is inhibitory/glutamatergic, so its effects on downstream partners are similar to those of an ON excitatory cell. (Note that information about whether synapses are excitatory or inhibitory was not used by our clustering algorithm.) Cluster9 also contains C2 and C3, which are expected to be ON cells because their top inputs are L1 and L5.

The ON and OFF motion pathways were traditionally defined by working backwards from the T4 and T5 motion detectors, which respectively compute the directions of moving ON and OFF stimuli (Borst and Groschner 2023). The ON motion pathway is directly upstream from T4 and includes Mi1, Mi4, Mi9, and Tm3. The OFF motion pathway is directly upstream from T5 and includes Tm1, Tm2, Tm4, and Tm9. These types are the consensus (Currier, Pang, and Clandinin 2023), though some authors include further types (Borst and Groschner 2023). The ON and OFF channels defined by our clustering have some overlap with the ON and OFF motion pathways, but they are not the same.

Interestingly, L3 does not belong to either Cluster8 or Cluster9. L3 connectivity is sufficiently unique that it stands apart from all other cell types as a cluster containing only the single type L3. This is consistent with current thinking that L3 constitutes a separate luminance channel, distinct from ON and OFF channels (Ketkar et al. 2022). L3 is the only L type with a sustained rather than transient response (Drews et al. 2020), and it encodes luminance rather than contrast (Ketkar et al. 2020).

Cluster3 consists of Dm4, Dm12, Dm20, and Mi9, which all have L3 as their strongest input. Mi9 is also the strongest output of L3, and like L3 exhibits a sustained response (Arenz et al. 2017). Therefore we include Cluster3 with L3 in the luminance channel. Mi9 is traditionally grouped in the ON motion pathway, but Mi9 is an input to the object and color subsystems, not only the motion subsystem.

The clusters also include less numerous types. The ON channel (Cluster9) contains interneuron types Dm1 and Dm18. A companion paper argues that the Dm1 and Dm8 normalize the activities of the numerous types in the ON pathway. The OFF channel (Cluster8) contains interneuron types from the Dm, Li, and Pm families.

### Motion

The motion-detecting T4 and T5 families belong to Cluster12. The cluster also contains CT1 and Tm9, which are well-known to be important for motion computation. It makes sense to regard Tm9 as dedicated to the motion subsystem, as the 80% of its output synapses are onto CT1 or T5. Tm9 should not belong to a general-purpose OFF channel. As a novel type, Li14 is the only surprise in Cluster12. Its strongest input is T5a, and its strongest outputs are T5a through T5d. T4/T5 neurons synapse onto VPNs that exit the optic lobe and enter the central brain (Data S4).

Cluster11 contains all of the lobula plate interneuron family, LPi1 through LPi15. Over half of these are new (Methods). Some LPi types consist of one or two cells that cover the entire visual field (Fig. 5b). Two LPi types may stratify in the same lobula plate layers, but consist of cells with different sizes (Fig. 5c). Most LPi types are amacrine, but some exhibit axo-dendritic polarization (Fig. 5d). Some types collectively cover only a portion of the visual field (e.g. LPi01 and LPi03 are ventral-only, Data S1).

**Figure 4.**
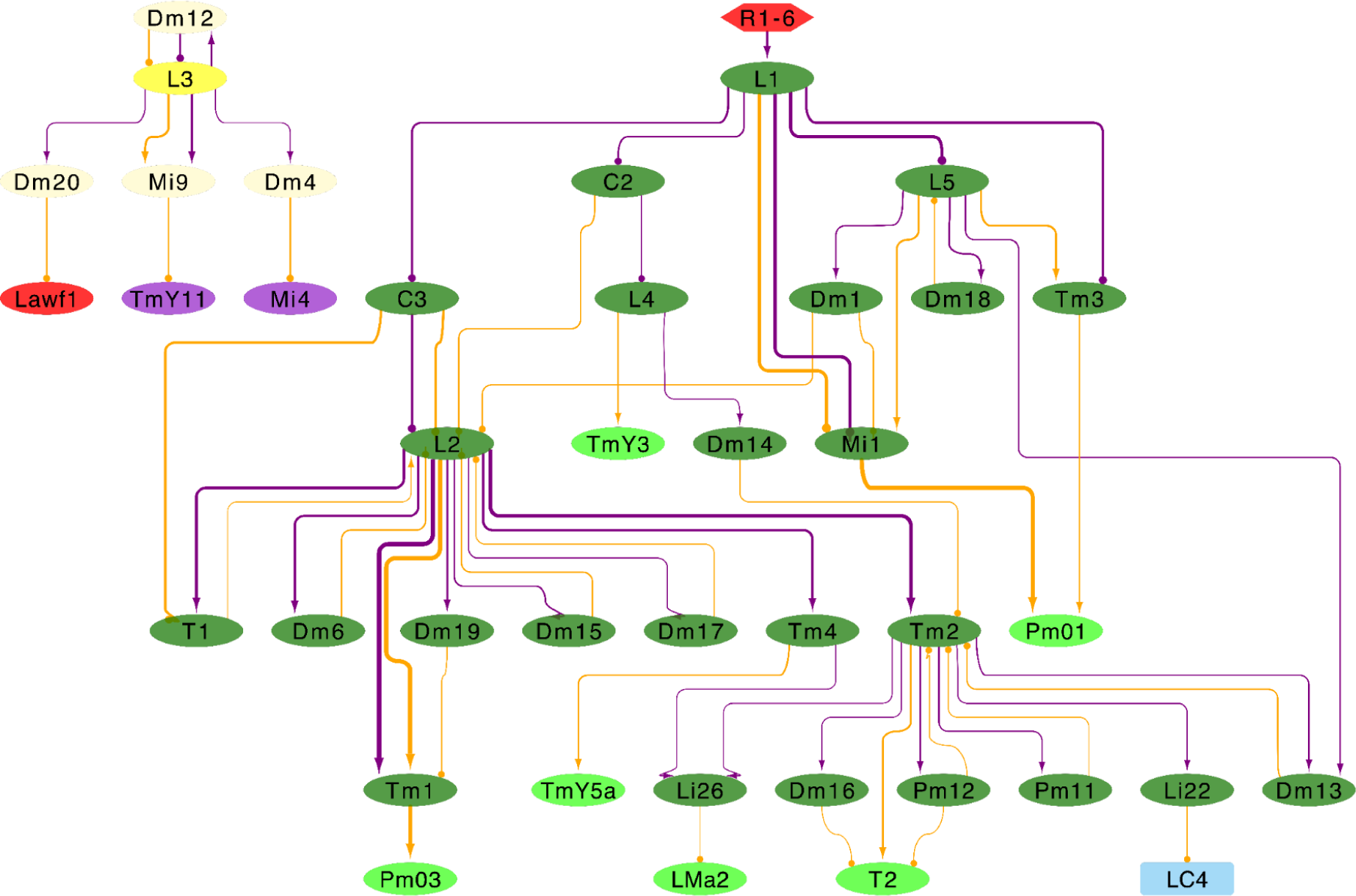
ON, OFF, and luminance channels. Simplified wiring diagram of ON, OFF, and luminance channels. Only strongest input and output connections are shown for clarity. Cell types (dark green) of the ON and OFF channels (Cluster9, Cluster8) are drawn with input and output types in other subsystems. Types (yellow, light yellow) in the luminance channel (L3 and Cluster3) are shown at upper left.

**Figure 5.**
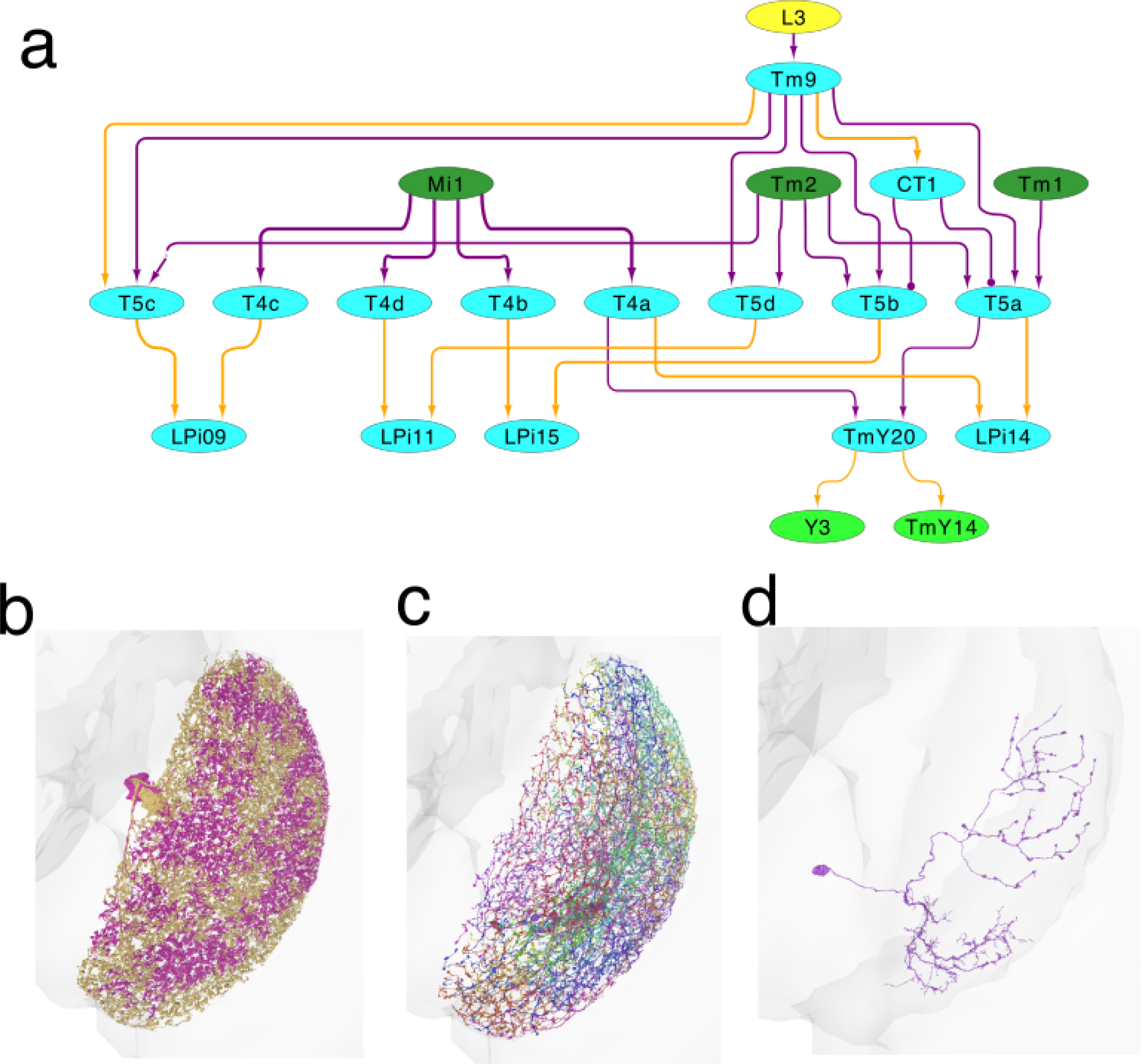
Motion subsystem. (a) Cell types (cyan) of the motion subsystem (Cluster11, Cluster12) containing more than 100 cells. Also shown are cell types from other subsystems that are connected to the motion subsystem. Only top input and output connections are shown for clarity. (b) LPi14 (LPi⇐T5a⇒H2), called LPi1-2 by (Shinomiya et al. 2022), is a jigsaw pair of full-field cells. (c) LPi02 (LPi⇐T5a⇒LPLC2) stratifies in the same lobula plate layers as LPi14, but the cells are smaller. (d) LPi08 (LPi⇐T5c⇒LPLC4) is an example of an interneuron that is not amacrine. It is polarized, with a bouton-bearing axon that is dorsally located relative to the dendrite.

All LPi types receive input from T4/T5 types, so it is clear that Cluster11 is related to motion vision. All LPi types receive input from T4/T5 cells with a single preferred direction (Fig. 5, Data S4). The only exception is LPi07, which receives inputs from T4/T5 cells with preferred directions c and d. LPi types synapse onto other LPi types and onto VPNs (Data S4).

Cluster11 also contains columnar neurons from three Y types and all Tlp types. All of these are predicted glutamatergic, and are reciprocally connected with T4/T5 of particular preferred directions. The only exception is Tlp5, which only receives input from T4a/T5a. The Y and Tlp types also connect with LPi and columnar VPN types (Shinomiya et al. 2022). TmY20 and Am1 also belong to Cluster11, and were previously identified as motion-related by (Shinomiya et al. 2022).

### Objects

Cluster7 includes the numerous types T2, T2a, and T3, which have been implicated in the detection of small objects (Keleş et al. 2020). Their downstream VPN partners are also involved in object detection. LC11 is activated by small objects (Keleş et al. 2020), and LC18 by very small objects (Klapoetke et al. 2022). LC17 is activated by small objects (Turner et al. 2022) and looming stimuli (Klapoetke et al. 2022). Based on this information, we will regard Cluster7 as part of the object subsystem.

Mi1 and Tm1 are the most prominent inputs to the object subsystem (Fig. 6), and respectively belong to the ON and OFF channels defined above. In spite of their names, T2a and T3 are more similar in their connectivity than they are to T2 (Fig. 2c). Mi1 and Tm1 are their top inputs, which explains why T2a and T3 are both ON-OFF (Keleş et al. 2020). T2 is ON-OFF because its top inputs are L5 and Tm2, which respectively belong to the ON and OFF channels.

**Figure 6.**
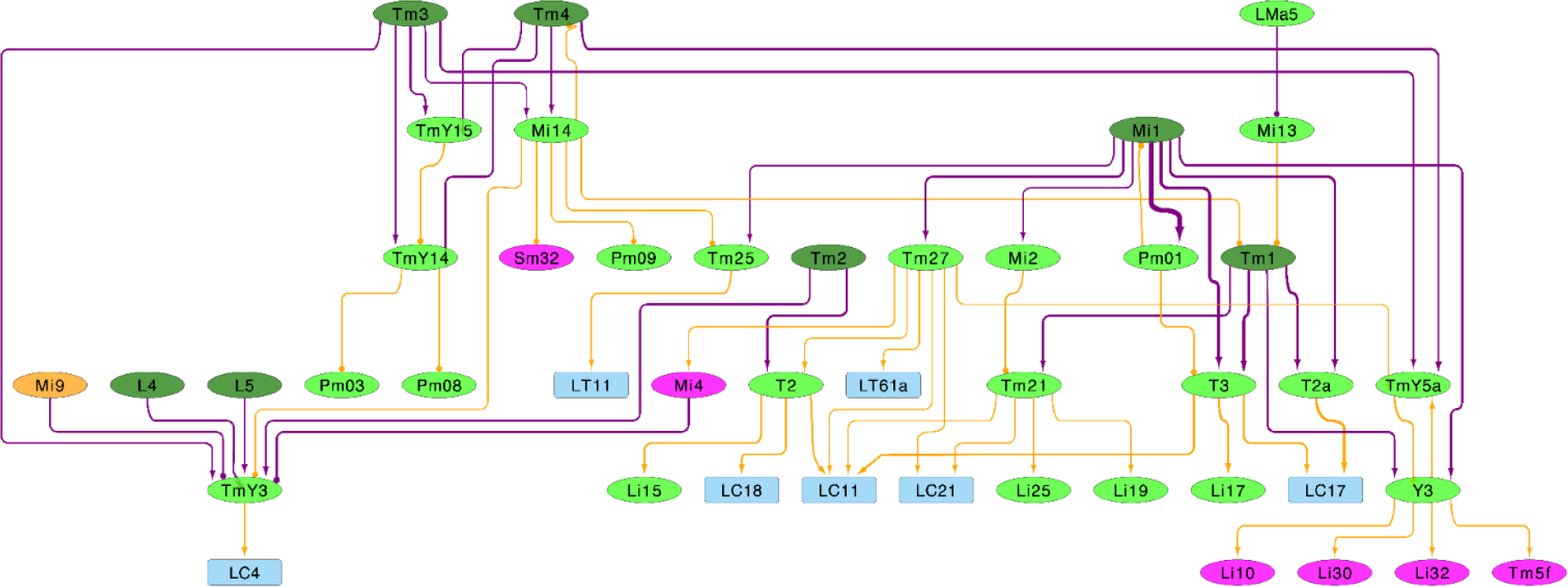
Object subsystem. Cell types (light green) of the object subsystem (Cluster7 and Cluster10) connect with VPNs (light blue), the color subsystem (magenta), and ON and OFF channels (dark green). Only top input and output connections are drawn for clarity.

Several types are nearby T2, T2a, and T3 in the dendrogram of the object subsystem (Fig. 2c). In particular, Tm21, Tm25, Tm27, TmY3, and Y3 are fairly numerous and excitatory, so we regard them as candidate object detectors. The object subsystem includes many other types from columnar families (Mi, TmY, Y), interneuron families (Li and Pm), and cross-neuropil tangential and amacrine families (LMa, LMt, MLt). Downstream targets of the object subsystem include LC, LPLC, and LT types (Fig. 6).

Cluster10 contains Li19 and Li25. The top input and output partner of both Li types is Tm21, which belongs to Cluster7. Therefore we include Cluster10 as well as Cluster7 in the object subsystem.

### Color and polarization

The inner photoreceptors R7 and R8 are important for *Drosophila* color vision, because their responses are more narrowly tuned to the wavelength of light than those of the outer photoreceptors R1-6. R7 prefers UV light, while R8 prefers blue or green light (Schnaitmann, Pagni, and Reiff 2020). We were able to proofread about 650 cells each for R7 and R8, a large fraction of the expected 800 cells. The automated synapse detector is unreliable for photoreceptors, as it was not trained to handle their dark cytoplasm. Therefore R7 and R8 connectivity onto postsynaptic targets is only qualitative in v783. A new synapse detector has been trained and is being applied to the data; the results will become available in the next release. R7 and R8 are contained in Cluster6, which also includes their targets Dm8, Dm9, Dm11, and DmDRA2 (Kind et al. 2021).

Cluster1 contains most of the remaining types so far implicated in color vision. As originally defined by (Fischbach and Dittrich 1989), Tm5 is a potential postsynaptic target of the inner photoreceptors, because it stratifies in the distal medulla at the M7 border and also in M3. These are the medulla layers containing the axon terminals of R7 and R8 (Gao et al. 2008). We found that Tm5 consists of six cell types (Fig. 7a). Three of our connectomic Tm5 types correspond to canonical Tm5 types that were previously defined by morphology and Ort expression (Gao et al. 2008; Karuppudurai et al. 2014). Tm5a and Tm5b receive R7 input, while Tm5c receives R8 input. In addition, we found three new types Tm5d, Tm5e, and Tm5f that receive little or no photoreceptor input, although their stratifications are similar to those of the canonical Tm5 types (Fig. 7a).

**Figure 7.**
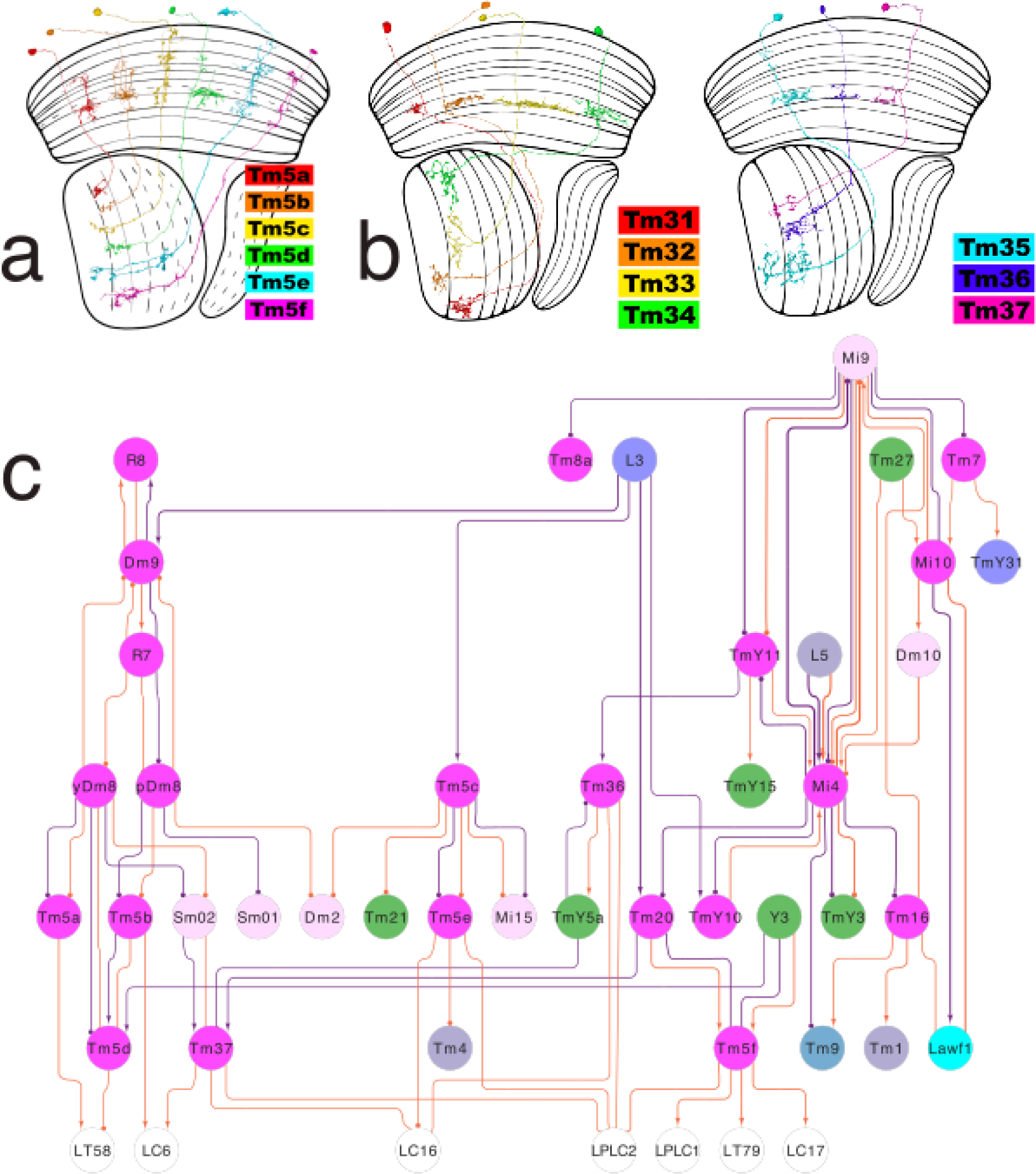
Color subsystem. (a) Tm5a through Tm5c correspond with types that were previously defined by molecular means. Tm5d through Tm5f have similar morphologies, but different connectivity patterns (Data S4). (b) Tm31 through Tm37 are new members of the Tm family that project from the serpentine layer (M7) to the lobula. (c) Cell types (magenta, pink) in the color subsystem (Cluster1, Cluster4, and Cluster6) that contain more than XX cells. Also shown are cell types that are inputs and outputs of the color subsystem. Only top input and output connections are shown for clarity.

The correspondences between connectomic and morphological-molecular Tm5 types were established using morphological criteria (Methods). However, the reader should be cautioned that there is considerable variability within a type, so reliably typing individual cells based on morphology alone is difficult or impossible. Connectivity is essential for reliable discriminations.

Tm5a and Tm5b receive R7 and Dm8 input, as expected from previous reports (Karuppudurai et al. 2014; Menon et al. 2019; Kind et al. 2021). Tm5c receives R8 input (Karuppudurai et al. 2014; Kind et al. 2021), and also strong L3 input (Fig. 7c, Data S4). While some synapses from Dm8 to Tm5c do exist (Karuppudurai et al. 2014), this connection is weak.

Tm20 has been implicated in color vision because it receives R8 input (S.-Y. Takemura et al. 2013, 2015; Kind et al. 2021). It also receives strong L3 input (Fig. 7b). Therefore Tm20 inputs are similar to Tm5c inputs, consistent with the physiological finding that these two types are more similar to each other in their chromatic responses than they are to Tm5a and Tm5b (Christenson et al. 2023).

Since Tm5a, Tm5b, Tm5c, and Tm20 are known to be related to color vision, we propose that the rest of Cluster1 is also part of a color subsystem. The new Tm5 types (Tm5d, e, and f) receive little or no synapses from photoreceptors, but they do receive indirect R7 and R8 input. Tm5d and Tm5e are predicted glutamatergic and Tm5f is predicted cholinergic. Tm5d receives indirect R7 input from Tm5b and Dm8a, Tm5e receives indirect R8 input from Tm5c, and Tm5f receives indirect R8 input from Tm20 (Fig. 7c).

We have defined Dm8a and Dm8b, which connect with Tm5a and Tm5b, respectively, and this preference is highly selective. As with Tm5, splitting Dm8 is straightforward with connectivity but difficult or impossible with morphology. Molecular studies previously divided Dm8 cells into two types (yDm8 and pDm8), depending on whether or not they express DIP*γ* (Menon et al. 2019; Courgeon and Desplan 2019). Physiological studies demonstrated that yDm8 and pDm8 have differing spectral sensitivities (Li et al. 2021). How yDm8 and pDm8 correspond with our Dm8a and Dm8b remains to be settled. Some preliminary clues are described in the Methods, but a definitive answer will require annotating yellow and pale columns using an improved version of photoreceptor synapses that is forthcoming.

Cluster1 also includes Tm7, Tm8a and Tm8b (another novel split), Tm16, and wholly new types Tm31 through Tm37. The latter deviate from the classical definition of the Tm family, which is supposed to project from the distal medulla to the lobula (Fischbach and Dittrich 1989). These types mainly stratify in the serpentine medulla (M7) and lobula, with little or no presence in the distal medulla (Fig. 7b). Nevertheless, we decided to lump them into the Tm family. Tm31 to Tm35 each contain relatively few (<100) cells, and are predicted not cholinergic. This departs from the norm for existing Tm types, which are generally more numerous (>100 cells) and predicted cholinergic. (Exceptions are the three glutamatergic Tm5 types). Tm36 and Tm37 contain more than 100 cells each, and are predicted to be cholinergic.

Cluster1 includes several TmY types, many Li and Sm interneuron types, two Pm interneuron types, some MLt types, and LLPt. Cluster1 also includes Mi4 and Mi10. Mi4 was traditionally regarded as part of the ON motion pathway, but makes more synapses onto types in the color subsystem than the motion subsystem. Its strongest output by far is Mi9, which we have assigned to a luminance channel and is one of the major inputs to the color subsystem. Mi4 also has output onto the object subsystem. This diversity of targets shows that Mi4 is a major hub, though our clustering has assigned it to the color subsystem. Mi10 mediates a feedback loop L3→Mi9→Mi10→Lawf1→L3, so it might seem to belong to the luminance channel, but it also has a number of partners in Cluster1.

Besides L3, Mi9 is another prominent input to the color subsystem (Fig. 7c). Both L3 and Mi9 belong to the luminance channel defined above. It makes sense that luminance information should be necessary for color computations (Ketkar et al. 2022).

Cluster4 contains Mi15 and Dm2, which are nearest neighbors in the dendrogram. Both types are known to receive inner photoreceptor input (Kind et al. 2021), so we also regard Cluster4 as part of the color subsystem. Cluster4 also contains many Sm types, two MLt, and Dm10. Dm2 is a major hub (Fig. 3), providing input to most Sm types (Data S4).

Cluster5 contains DmDRA1, a cell type at the dorsal rim of the medulla known to be important for behaviors that depend on skylight polarization (Sancer et al. 2019). Cluster5 is therefore regarded as part of the polarization subsystem. It contains several Sm types, most of which are either situated at the dorsal rim or have some specialization there. DmDRA2 should also be part of the polarization subsystem, but is currently assigned to the color-related Cluster6. That may be because we do not currently distinguish R7 and R8 in the dorsal rim area from other R7 and R8.

### Morphological variation

As mentioned earlier, connectivity is important for distinguishing between types with similar morphologies. Connectivity is also helpful for deciding when to ignore morphological variations between cells of the same type. A common kind of variation involves the targeting of the axon. For example, TmY14 was originally identified by (S.-Y. Takemura et al. 2013) as a cell type intrinsic to the optic lobe, but (Shinomiya et al. 2019) later recognized that TmY14 might be classified as a VPN, because it projects to the central brain. In another story twist, our optic lobe turns out to contain a subset of TmY14 that lacks the central brain projection (Fig. 8a, b). In these atypical TmY14 cells, the axon remains in the medulla rather than projecting to the central brain. Whether typical or atypical, the TmY14 axon has few synapses and minimal impact on connectivity. And the optic lobe connectivity of the TmY14 cells seems not to depend on whether or not there is a central brain projection. Therefore we have ignored the inconsistent axon, and regard TmY14 as a single type that is intrinsic to the optic lobe. (In cases like this, we double check the proofreading before concluding that this is true morphological variation rather than an experimental artifact.)

**Figure 8.**
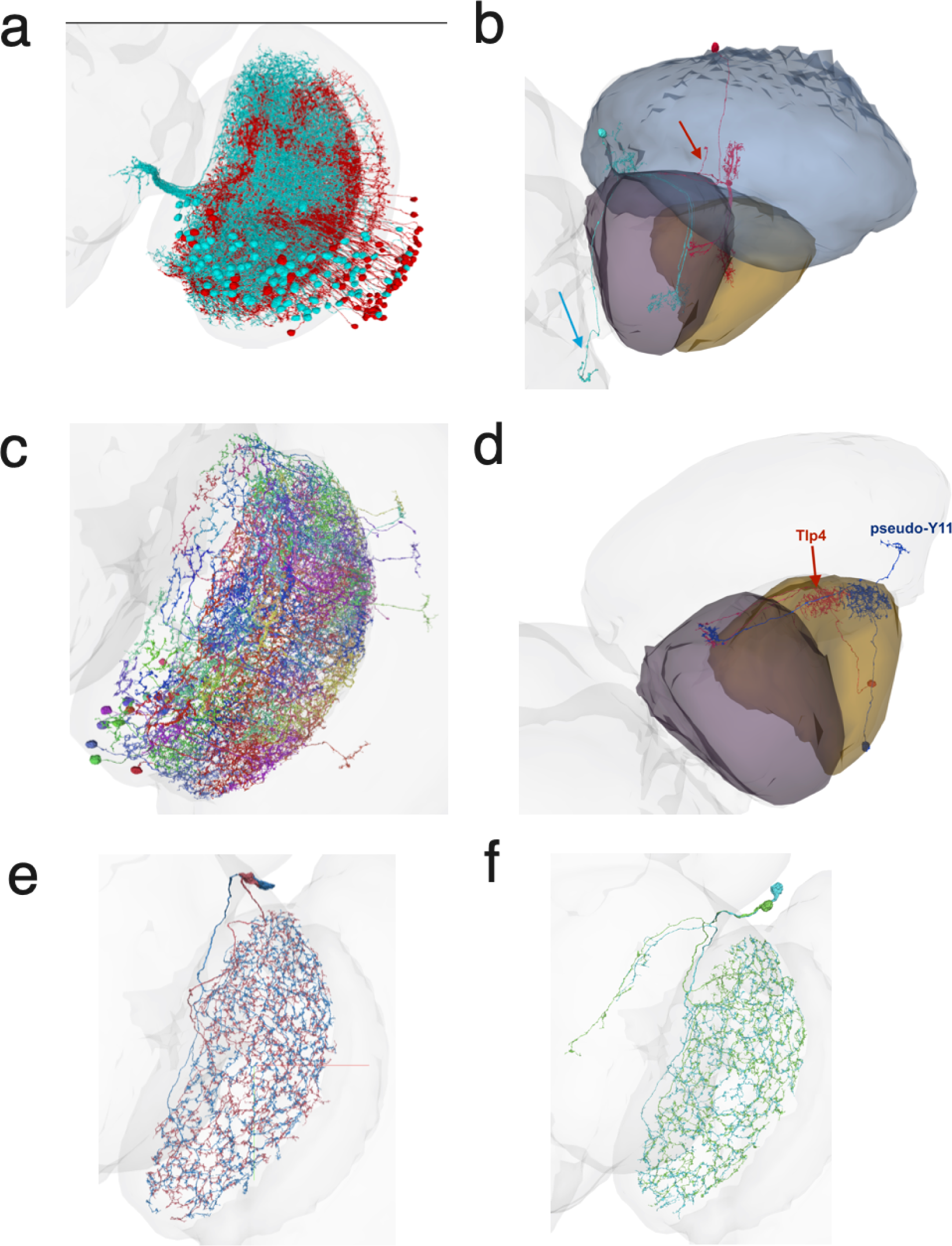
Morphological variation. (a) The TmY14 cell type exhibits a mixture of typical and atypical cell morphologies, with projections extending to the central brain or the medulla. Some of the atypical ones project toward the central brain but retract without reaching it, instead projecting into the medulla. Their connectivity remains consistent regardless of whether the cell projects to the central brain. (b) Representative typical (cyan) and atypical (red) TmY14 with arbor projecting into the central brain and medulla respectively. (c) A few Tlp4 cells exhibit Y11 - like morphology, but have the same connectivity as Tlp4. We call these cells pseudo-Y11. (d) Morphological comparison of Tlp4 and pseudo-Y11. Pseudo-Y11 has an additional branch in the medulla. (e) Li11 does not project into the central brain. (f) Pseudo-Li11 has an additional arbor projection into the central brain. This arbor makes a few synapses, and might lead to the conclusion that pseudo-Li11 should be categorized as Li11. However, the connectivity between Li11 and pseudo-Li11 is fundamentally different, making them distinct types.

Another interesting example is Tlp4 versus Y11. These two types have similar connectivity patterns (Data S4), and are near each other in the hierarchical clustering of types (Fig. 2c). A major difference is that Tlp4 cells, by definition, have no connectivity in the medulla. Except a few of them do, and look like they do not belong in Tlp4 (Fig. 8d). For a long time, we classified these errant cells as Y11. But it turns out that their medullar projections make few synapses, and their connectivity in the lobula and lobula plate matches Tlp4. Therefore we have included such “pseudo-Y11” cells in the Tlp4 type.

It is worth mentioning an unusual example in which ignoring morphological variation is correct in one sense, but ultimately turns out to be misleading. Three Li11 cells are annotated in the hemibrain (Scheffer et al. 2020), and three corresponding cells can be identified in our optic lobe (Schlegel et al. 2023). We group two of these cells in one type (Fig. 8e). The third cell can be paired with a fourth to form a “pseudo-Li11” type with a small projection into the central brain (Fig. 8f). This is a striking difference between Li11 and pseudo-Li11 morphologies, but the central brain projection has few synapses and therefore little impact on connectivity. Therefore it might be tempting to ignore the projection as a developmental “accident” and merge Li11 and pseudo-Li11 into a single type. But it turns out that Li11 and pseudo-Li11 can be distinguished by connectivity in the lobula. For example, Li25 has strong LT61 output, while Li19 has strong LT11 input. Pseudo-Li11 also exists in the hemibrain (Schlegel et al. personal communication), though there it lacks the small projection. So the central brain projection of pseudo-Li11 exhibits variability across individuals, further evidence that it is a developmental accident. Rather than Li11 and pseudo-Li11, we use the names Li25 (Li⇐Tm21⇒LT61) and Li19 (Li⇐Tm21⇐LT11).

We also saw a few “weirdo” cells, which looked strange and were usually one-of-a-kind. For example, cell 720575940629614953 resembles an Li full-field cell, but also extends a smaller secondary arbor into the lobula plate and medulla. Originally we decided that this was a developmental accident, and did not include it in our list of types. More recently, we found that this odd-looking cell is repeated in the left optic lobe, and have promoted it to a type (Li29).

### Spatial coverage

Our description of the connectomic approach to cell typing glossed over an important issue. Whether to split types more finely or merge types more coarsely may seem subjective. Fortunately, we were able to resolve this lumper-splitter dilemma by using spatial coverage as a criterion.

As a general rule, the cells of a cell type collectively cover all columns of the optic lobe with a density that is fairly uniform across the visual field. This makes sense for implementing translation-invariant computations at every location in the visual field, a strategy commonly used in convolutional networks and other computer vision algorithms. Uniform spatial coverage is sometimes called “tiling,” but for the less numerous types the arbors typically overlap so much that the analogy to floor tiles is misleading. Spatial coverage is also a property of many cell types in mammalian retina (H. Sebastian Seung and Sümbül 2014; Bae et al. 2018).

In some types consisting of just one or a few cells, we discovered an unconventional “jigsaw” style spatial coverage. For example, LPi14 (LPi⇐T5a⇒H2), called LPi1-2 by (Shinomiya et al. 2022), is a pair of full-field cells (Fig. 5b). We refer to them as a “jigsaw pair” because they jointly cover the visual field in an irregular fashion, as if they were cut by a jigsaw. Jigsaw types can also be found in other interneuron families and include Pm14, Li27, and Li28.

Our feature vector (Fig. 2a) includes no explicit information about the spatial coordinates of a cell. Therefore, if clustering feature vectors results in cell types with good spatial coverage, that is a validation of the clustering. Coverage also solves the lumper-splitter dilemma. Suppose we attempt to split one type into two candidate types, based on hierarchical clustering. If both candidate types exhibit good coverage, then we accept them as valid. If the cells of both candidate types seem randomly scattered, that means our split is invalid, because it is presumably discriminating between cells based on noise. (Chromatic types might seem like an exception to this rule, but their locations are only apparently random because they depend on the locations of pale and yellow columns.)

The above are easy cases, but there are also edge cases. Suppose that the two candidate types cover the dorsal field and the ventral field, respectively. In this case, we reject the split, and prefer to say that there is a single type that exhibits dorsoventral spatial variation in connectivity. On the other hand, if one candidate type covers the dorsal field and the other covers the full field, this is an acceptable split. Conversely, if hierarchical clustering merges two types with nonoverlapping spatial coverage, the result is likely a correct type. If hierarchical clustering merges two types with overlapping coverage, the result is likely not correct.

With these heuristics, some of our cell types end up with only partial coverage of the visual field (Fig. 9). This is especially common for boundary types. Sm is the intrinsic type family containing the most types with partial coverage. This makes sense, given that Sm cells interact closely with many boundary types arborizing in the serpentine layer. Cell types with partial coverage make sense in the later stages of vision. After the early stages of vision, computer vision also often discards translation invariance and may perform different visual computations in different regions of the visual field.

**Figure 9.**
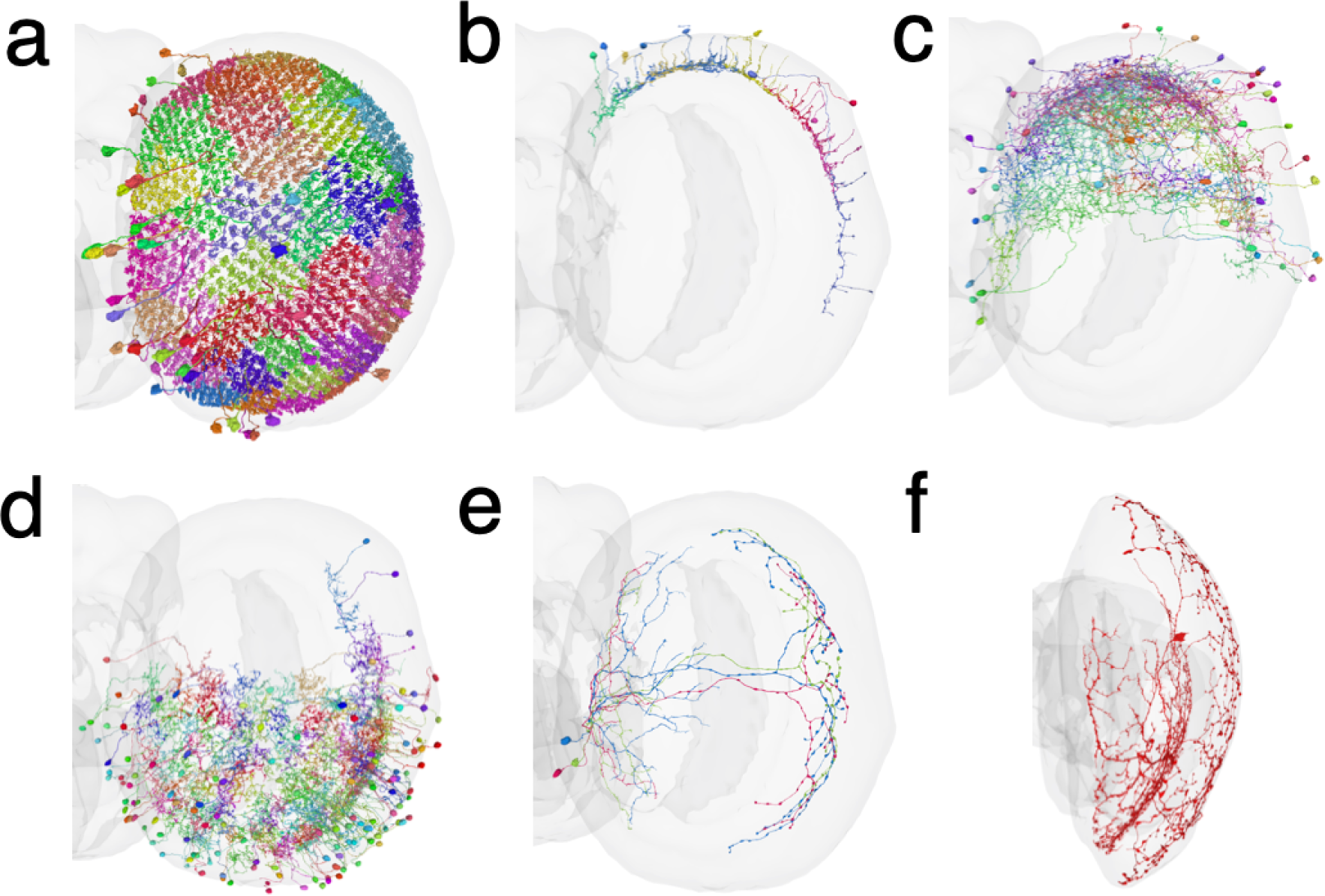
Different kinds of spatial coverage. (a) Example (Dm4) of complete tiling with no overlap (b) Example (DmDRA2) of dorsal rim coverage (c) Example (Sm12, Sm⇐MC65⇒TmY5a) of dorsal hemifield coverage (d) Example (Sm01, Sm⇐Mi9⇒CB0165) of ventral hemifield coverage (e) Example (Sm33, Sm⇐IB029⇐MeTu1) of H-shaped coverage (anterior and posterior rim) (f) Singleton (Sm39, Sm⇐aMe4⇒Mi15) with mixed coverage: dorsal dendritic arbor in M7 and full-field axonal arbor in M1.

## Discussion

The connectomic approach to cell typing has three powers. First, it does not suffer from the incomplete and biased sampling that can plague other methods. Second, connectivity turns out to provide a rich set of features for distinguishing between cell types. Third, connectomic cell typing not only yields cell types, but even more importantly tells us how they are wired to each other.

### Implications for visual function

We clustered cell types with similar connectivity patterns (Fig. 2c), and the resulting clusters turned out to be interpretable in terms of visual functions. We were able to define subsystems for motion, object, and color vision (Figs. 5, 6, 7), which are fed by ON, OFF, and luminance channels (Fig. 4).

The motion subsystem (Cluster12, Cluster11) contains 35 types. Of the 11 types in Cluster12, eight are the well-studied T4/T5 types, and only one (Li14) has never yet been recorded by physiologists. Less is known about Cluster11; of its 24 types, only LPi09 (LPi3-4) and LPi11 (LPi4-3) have been recorded by physiologists (Mauss et al. 2015). It has been proposed that LPi types mediate opponent interactions between cells that are activated by different directions of motion (Mauss et al. 2015). (Ammer et al. 2023) provided evidence for such opponency using the particular pair of LPi types, LPi09 (LPi3-4) and LPi11 (LPi4-3). Our data can be used to test the generality of their hypothesis for all LPi types, over half of which are new. It turns out that there are some exceptions to opponency, as will be described in a companion paper.

Of the 49 types in the object subsystem (Cluster7, Cluster10), T2, T2a, and T3 have been characterized by physiologists as object detectors (Keleş et al. 2020). Our work suggests that the remaining types are also involved in object detection.

The color subsystem (Cluster6, Cluster1, Cluster4) contains almost 100 types, almost half of the intrinsic types in the optic lobe. One can only speculate about the reason for this numeric preponderance. Perhaps color vision is especially important for the fly, or an especially complex computation. Or perhaps color vision is an overly simplified name for what the color subsystem actually does. More than half of the color types are Sm and Li interneurons.

We have neglected Cluster2 (Fig. 2c), which a companion paper argues is a subsystem for form vision (H. Sebastian Seung 2023). The form subsystem contains just six types that are predicted to compute the orientations of visual stimuli. The small number of cell types suggests that form vision might be less important and/or less well-developed in *Drosophila* than in mammalian visual systems.

We have carved the optic lobe into distinct subsystems, but there are also interactions between the subsystems. Perusal of the connectivity data readily reveals “crosstalk” connections between cell types in different subsystems (Data S4).

### Implications for visual development

Mammalian visual cortex was historically fertile ground for studies of neural circuit development, starting with attempts to explain ocular dominance and orientation columns (Katz and Crowley 2002). These aspects of cortical organization can be seen without single cell resolution, and now seem crude compared to neuronal wiring diagrams. Now that we have a detailed wiring diagram for the fly visual system, we can regard it as the end goal of fly visual system development. Single cell transcriptomics is providing detailed information about the molecules in fly visual neurons (Kurmangaliyev et al. 2020; Özel et al. 2021; Konstantinides et al. 2022). Comparison of transcriptomic and connectomic information is already uncovering molecules important for the development of the fly visual system (Yoo et al. 2023), and this trend is bound to increase in momentum.

### Complete and unbiased

Most of our new types are not so common (10 to 100 cells), which may be why they were missed by (Fischbach and Dittrich 1989). These pioneers relied on Golgi staining to sample neurons from multiple individuals, a technique that is well-suited for identifying the most numerous types. We can only speculate about why they missed the Sm family entirely. And we do not know why so many types defined by (Fischbach and Dittrich 1989) cannot be identified in our optic lobe. These authors acknowledge that Golgi staining yields a biased sample of cells, and that morphological variability might have caused them to overestimate the number of types.

Contemporary light microscopic anatomy is more powerful because it can leverage the selective labeling of genetic lines. However, combining light microscopy with genetic control does not evade the limitations of incomplete and biased sampling. The story of Tm5 serves as a case in point. A breakthrough in color vision started by genetically labeling neurons that express the histamine receptor Ort (Gao et al. 2008). Researchers reasoned Ort would be expressed by cells postsynaptic to the chromatic photoreceptors R7 and R8, which are histaminergic. Then light microscopic anatomy was used to make fine distinctions between three Tm5 types labeled in the transgenic line (Gao et al. 2008). The present connectomic work has revealed six Tm5 types, a finding that was only foreshadowed by previous work on the same EM dataset (Kind et al. 2021). The three new Tm5 types were presumably missed by previous work because they receive little or no direct photoreceptor input (Fig. 7b), and do not express Ort. Nevertheless, they belong to the color subsystem because they are strongly interconnected with types that receive direct R7 and R8 input. For example, Tm5d might be hue selective, because its strongest input is Tm5b, which is known to be hue selective (Christenson et al. 2023).

The Tm5 example demonstrates that connectomics can find fresh patches in well-trodden ground. More telling is that connectomics can guide us to entirely new landscapes, such as the 42 Sm types in an entirely new type family.

### Distinguishing cell types using connectivity

The first and second stages of our cell typing relied chiefly on single-cell anatomy (Methods). The third stage, however, had to rely heavily on connectivity. Connectivity-based features (Fig. 2a) enabled us to discriminate between cell types that stratify in very similar neuropil layers. This was essential, for example, to define Sm types. They are easy to confuse because they are so thinly stratified at or near the serpentine layer (M7).

Stratification constrains connectivity, because neurons cannot connect with each other unless they overlap in the same layers (Masland 2004). However, stratification does not completely determine connectivity, because neurons in the same layer may or may not connect with each other. Classical neuroanatomy, whether based on Golgi or genetic staining, relied on stratification because it could be seen with a light microscope. Now that we have electron microscopic data, we can rely on connectivity for cell typing, rather than settle for stratification and other properties from single-cell anatomy (H. Sebastian Seung and Sümbül 2014).

We have additionally verified self-consistency of the final types in the connectivity feature space. Virtually every cell is assigned to the nearest type in feature space, where nearest is defined by Jaccard distance. Such self-consistency might be viewed as trivial because erroneous type assignments were corrected using Jaccard distance in the final stage of cell typing (see Clustering of high dimensional feature vectors and Methods). It is actually nontrivial because human experts have other means of type assignment beyond the feature vector. They can look at morphology, how the cells tile the visual field, and so on. The nontrivial result is that our type assignments are both consistent with expert opinion and self-consistent according to distances in feature space.

As noted previously (Scheffer et al. 2020), using this feature vector to define types is circular because the feature vector itself depends on the assignment of cells to types. In the initial stage of cell typing, these assignments are based on morphology as well as connectivity. Once the feature vector has become rich enough based on these assignments, it becomes possible to switch to clustering based on connectivity alone. We leave for future work the challenge of evolving this into a connectivity-based clustering algorithm from start to finish.

### Role of boundary types

If we include the almost 500 boundary types (Schlegel et al. 2023), there are 700+ cell types in the optic lobe. This is considerably greater than transcriptomic estimates of neuronal diversity. For example, one study reported 171 neuronal cell types, a figure that includes VPNs as well as intrinsic neurons (Özel et al. 2021).

Some might not be surprised by our empirical finding that defining intrinsic types can be done using only synapses between intrinsic neurons, because these make up 80% of all synapses of intrinsic neurons. Others might be skeptical of our empirical finding because boundary types contribute 20% of the synapses of intrinsic neurons, a fraction that may not sound insignificant. To explain our empirical finding, we speculate that synapses between intrinsic and boundary neurons do not aid typing because the longer feature vector has more redundancy in the information theoretic sense.

While boundary types are not necessary for defining intrinsic types, they are important to our work because the connectivity between types is arguably the main product of our work, and connectivity between intrinsic and boundary types is of great importance for understanding visual function.

Numerous VPN types project from the lobula to the central brain. The many LC types receive inputs from the color and object subsystems, some inputs from the form subsystem, and little input from the motion subsystem.

### Spatial organization

The wiring diagram for cell types (Fig. 3, S4, S5, S6) is simpler than the connectome from which it was derived. Part of the reduction in complexity comes from ignoring space. If we say that cell types A and B are connected, it is understood implicitly that this rule only applies when the tangential separation between the cells is small enough that their arbors overlap. In some cases connectivity might also depend on the absolute locations of the cells, not only their relative location. Cell types that cover only parts of the visual field (Fig. 9) are the clearest example of dependence on absolute location. The dependence of connectivity on space is essential for vision, and we leave its characterization for future work. Cell type labels and spatial coordinates can be regarded as discrete and continuous latent variables in models of connectivity (H. Sebastian Seung 2009).

### Artificial intelligence

This paper began by recounting the story (H. S. Seung and Yuste 2010) of how wiring diagrams for visual cortex drawn in the 1960s inspired convolutional nets, which eventually sparked the deep learning revolution in artificial intelligence (AI). Convolutional nets have now been applied to reconstruct the fly brain from electron microscopic images (Dorkenwald, Matsliah, et al. 2023), making the current study possible. Coming full circle, the fly optic lobe turns out to be as literal an implementation of a convolutional net as one could ever expect from a biological system. The columns of the optic lobe form a hexagonal lattice, rather than the square lattice used in computer vision, but it is a highly regular lattice nonetheless. And the activities of the neurons in each cell type are analogous to a feature map in a convolutional net (Lappalainen et al. 2023). One difference is that optic lobe circuits are highly recurrent, which is not the norm for the convolutional nets used in computer vision.

Another difference is that the connections of optic lobe neurons do not appear to be learned in the sense of AI, because they seem to have little dependence on visual experience (Scott, Reuter, and Luo 2003). However, mechanisms based on spontaneous activity (rather than visually evoked activity) might play a role in *Drosophila* visual development (Akin and Zipursky 2020), analogous to mammalian visual development.

### Implications for mammalian cell types

*Drosophila* serves as an interesting middle ground between small nervous systems and big mammalian brains. In *C. elegans*, the number of cell types is somewhat less than half the number of neurons; most cell types consist of just a pair of mirror symmetric neurons (White et al. 1986). Over 30,000 neurons are intrinsic to the central brain (Fig. S1a) of *Drosophila*, 100× more than *C. elegans*. Many cell types in the central brain consist of just one or a few neurons per hemisphere (Scheffer et al. 2020; Schlegel et al. 2023), meaning that the number of cell types is in the thousands (Fig. S1e). In the optic lobes, the number of cell types is far less than the number of neurons. A large ratio of neurons to cell types is reminiscent of mammalian retina or cortex, where the ratio is even larger (Zeng and Sanes 2017; BRAIN Initiative Cell Census Network (BICCN) 2021).

For cortical cell types, single cell transcriptomics has been hailed as more advanced than the old-fashioned anatomical approach (Yuste et al. 2020). Proponents are now coming around to a more nuanced view, because neurons of the same transcriptomic type can have highly variable morphological and electrical properties (Scala et al. 2020; Gouwens et al. 2020). It is not yet clear why this is the case. One possibility has long been suggested, which is that transcriptomic differences might exist during development, and vanish in adulthood. This possibility has been confirmed in the fly visual system. For example, T4 and T5 types with a and b preferred directions can be transcriptionally distinguished from those with c and d preferred directions in adult flies (Özel et al. 2021). But all four preferred directions can be transcriptionally distinguished only at the P50 pupal stage.

The connectomic approach is already being applied to cell types in visual cortex (Schneider-Mizell et al. 2023). There is obvious motivation to scale up the approach, and make it as definitive for the cortex as it now is for the fly visual system. Here the limiting factor is throughput. In mammalian brains, transcriptomics has so far had higher throughput, measured in number of cells per unit of time or money. This is because basic research in transcriptomics can leverage technologies developed by the large DNA sequencing industry. Connectomics is also becoming more economical, and this trend should continue.

### Future releases

This manuscript is based on v783, which is currently accessible through the FlyWire Codex (codex.flywire.ai). Another update will occur after the next release of the connectome. One major change will be more accurate detection of synapses, especially those made by photoreceptors. More proofread cells will also become available in the next release, and are already available through CAVEclient (Dorkenwald, Matsliah, et al. 2023; Dorkenwald, Schneider-Mizell, et al. 2023).

### Data availability

Most up to date optic lobe intrinsic cell type annotations can be downloaded directly from the Codex download portal (https://codex.flywire.ai/api/download). They are also versioned and available to download from a dedicated data repository on GitHub: https://github.com/murthylab/flywire-visual-neuron-types

These annotations will be ported to the next FlyWire Connectome releases when published.

## The FlyWire Consortium (optic lobe)

Krzysztof Kruk^2^, Ben Silverman^1^, Jay Gager^1^, Kyle Patrick Willie^1^, Doug Bland^1^, Austin T Burke^1^, James Hebditch^1^, Ryan Willie^1^, Celia David^1^, Gizem Sancer^8^, Jenna Joroff^4^, Dustin Garner^5^, annkri (Anne Kristiansen)^2^, Thomas Stocks^2^, Amalia Braun^6^, Szi-chieh Yu^1^, Marion Silies^7^, AzureJay (Jaime Skelton)^2^, TR77^2^, Maria Ioannidou^7^, Marissa Sorek^1^, Matt Collie^4^, Gerit Linneweber^8^, Sebastian Mauricio Molina Obando^7^, Rey Adrian Candilada^1^, Alexander Borst^6^, Wei-Chung Lee^4^, Philipp Schlegel^9,10^, Greg Jefferis^9,10^, Arie Matsliah^1^, Amy R. Sterling^1^, Emil Kind^8^, Mathias Wernet^8^, Sung Soo Kim^5^, Mala Murthy^1^, H. Sebastian Seung^1,11^

^1^Princeton Neuroscience Institute, Princeton University, Princeton, USA

^2^Eyewire, Boston, USA

^3^Department of Neuroscience, Yale University, New Haven, USA

^4^Harvard Medical School, Boston, USA

^5^University of California, Santa Barbara, USA

^6^Department Circuits-Computation-Models, Max Planck Institute for Biological Intelligence, Planegg, Germany

^7^Johannes-Gutenberg University Mainz, Mainz, Germany

^8^Institut für Biologie - Neurobiologie, Freie Universität Berlin, Germany

^9^Drosophila Connectomics Group, Department of Zoology, University of Cambridge, Cambridge, UK;

^10^Neurobiology Division, MRC Laboratory of Molecular Biology, Cambridge, UK;

^11^Computer Science Dept, Princeton University, Princeton, USA

## Acknowledgements

We are indebted to all those who proofread optic lobe intrinsic neurons (Dorkenwald, Matsliah, et al. 2023). We thank A. Nern and M. Reiser for educating and advising FlyWire community members who engaged in annotation of visual neurons. We thank the FAFB tracing community for supportive and open sharing of methods and data, especially the FAFB optic lobe working group. We are grateful to Y. Kurmangaliyev for his thoughtful comments on the manuscript. We thank R. Behnia and M. Silies for feedback about visual motion detection pathways. We thank J. Wiggins, G. McGrath, and D. Barlieb for computer system administration and M. Husseini for project administration. We are grateful to J. Maitin-Shepard for Neuroglancer. MM and HSS acknowledge support from the National Institutes of Health (NIH) BRAIN Initiative RF1 MH117815, RF1 MH129268 and U24 NS126935, from the Princeton Neuroscience Institute, as well as assistance from Google. DG, GS, and SK were supported by the National Eye Institute of the NIH (DP2EY032737), Searle Scholars Program, Sloan Research Fellowship, and Klingenstein-Simons Fellowship in Neuroscience. EK and MW were supported by Deutsche Forschungsgemeinschaft (DFG) grant WE 5761/4-1, SPP 2205, FOR 5289, and AFOSR grant FA9550-19-1-7005.

## Author Contributions

DG and GS annotated cells under the supervision of SSK and MW. MS and ARS recruited, trained, and managed citizen scientists with help from EK. KK annotated cells and created computational cell typing tools for use by the community. SY trained and managed DB, AB, JG, JH, BS, KW, RW to annotate the remaining known cell types and discover and annotate new types. AM and HSS created semiautomated cell typing tools. AM and HSS carried out the final automated stage of typing. AM verified types with predicates. HSS verified types with 2D projections. SY, HSS, MM devised type family names and composite type names. AM and HSS characterized subsystems. SSK, MW and MM identified implications for visual function. AB, JG, JH, BS, KW, RW, SY, AM, HSS created figures. KK, MS, ARS, AM, SY, SSK, MM, HSS wrote the text. MM and HSS supervised the project.

## Supplementary Figures

**Figure S1.**
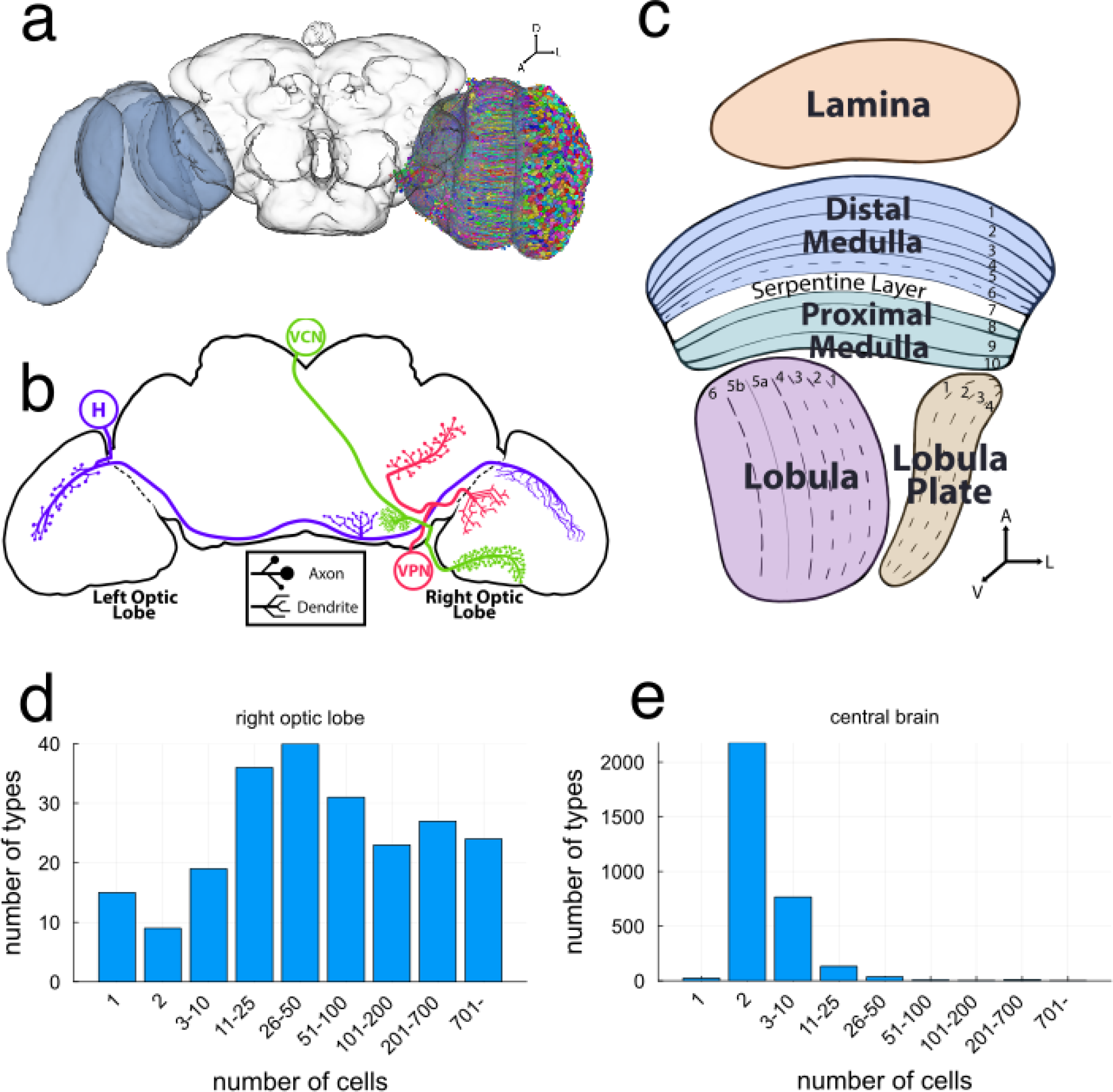
Cell counts of optic lobe types versus central brain types. (a) *Drosophila* central brain and flanking optic lobes. Neurons intrinsic to the right optic lobe (colors) are the subject of this study (A: Anterior. L: Lateral. D: Dorsal) (b) Boundary cells straddle the optic lobe and central brain (H: heterolateral, VCN: visual centrifugal neuron: VPN: visual projection neuron). (c) Optic lobe main neuropils (brain regions) and their layering. (A: Anterior. L: Lateral. V: Ventral) (d) Distribution of number of **optic lobe** types by bucketed unilateral cardinality. Each bar represents types whose cardinality (number of cells) is within the specified range. Most types contain 10+ cells, and a significant portion of types contain hundreds of cells. (e) Distribution of the number of **central brain** types by bucketed bilateral cardinality. In contrast to the optic lobe, here most types have cardinality 2 (cell and its mirror twin in the opposite hemisphere).

**Figure S2.**
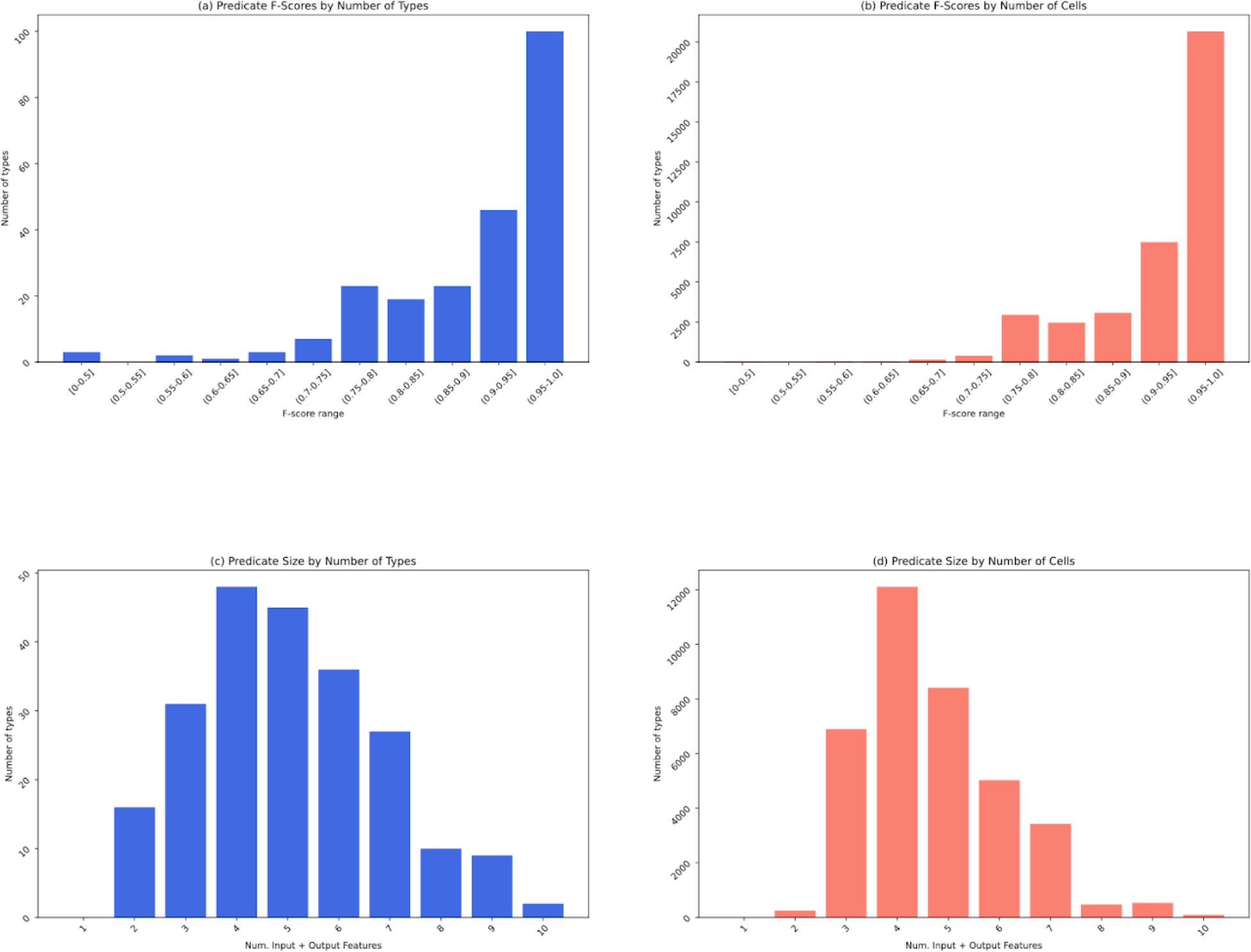
Logical connectivity predicate statistics. (a) Number of types by predicate f-score range. (b) Number of cells by their types’ predicate f-score range. (c) Number of types by predicate size, that is the sum of the number of input features and output features participating in the binary conjunction. (d) Number of cells by their types’ predicate size.

**Figure S3.**
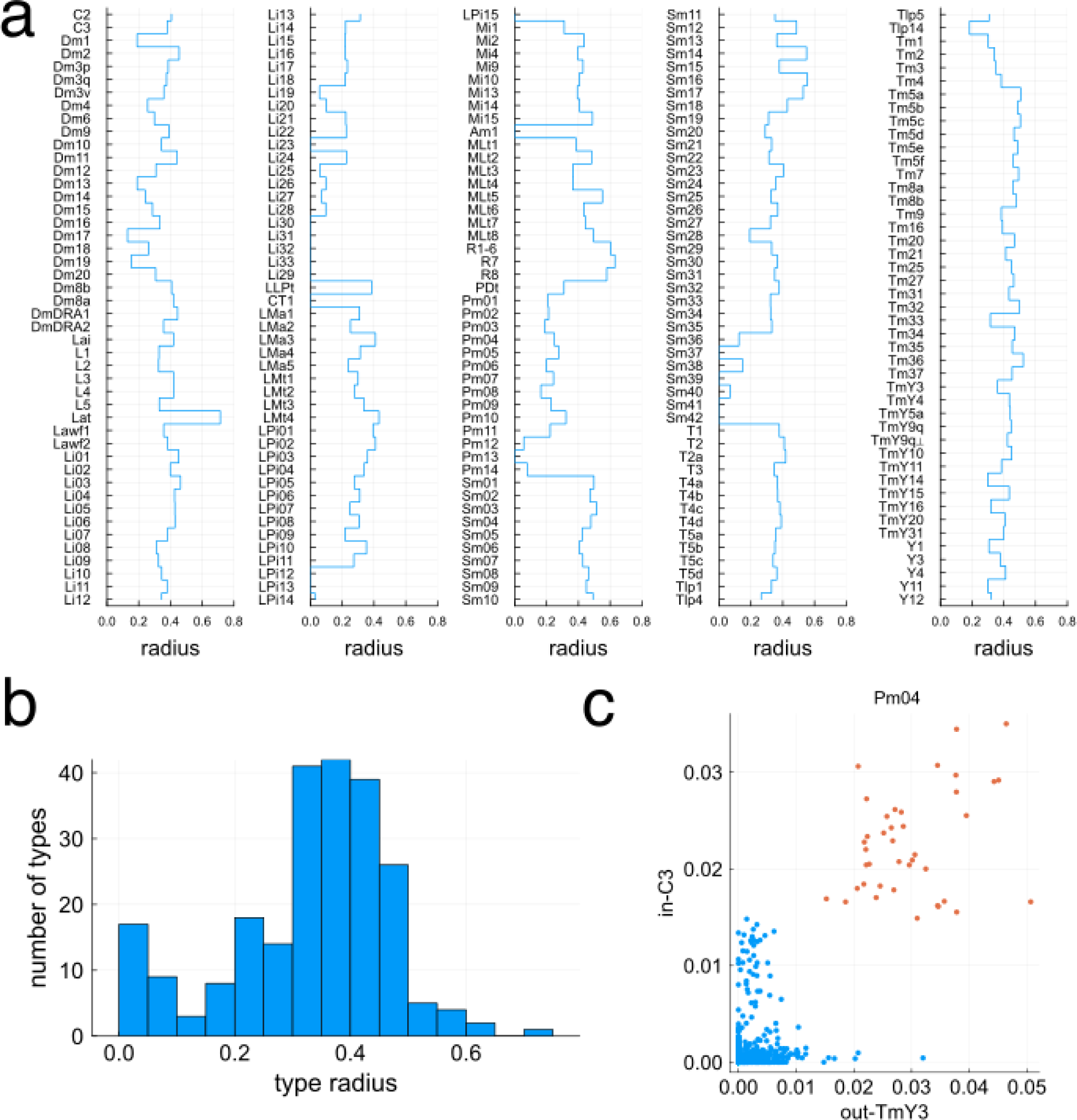
Discrimination in high and low dimensions. (a) Radii of types in high-dimensional feature space. (b) Histogram of type radii in high-dimensional feature space. (c) Example 2D discriminator for Pm04 cells (red) versus other Pm types (blue). On the X and Y axis are the fraction of their inputs / outputs in C3 / TmY3 respectively.

**Figure S4.**
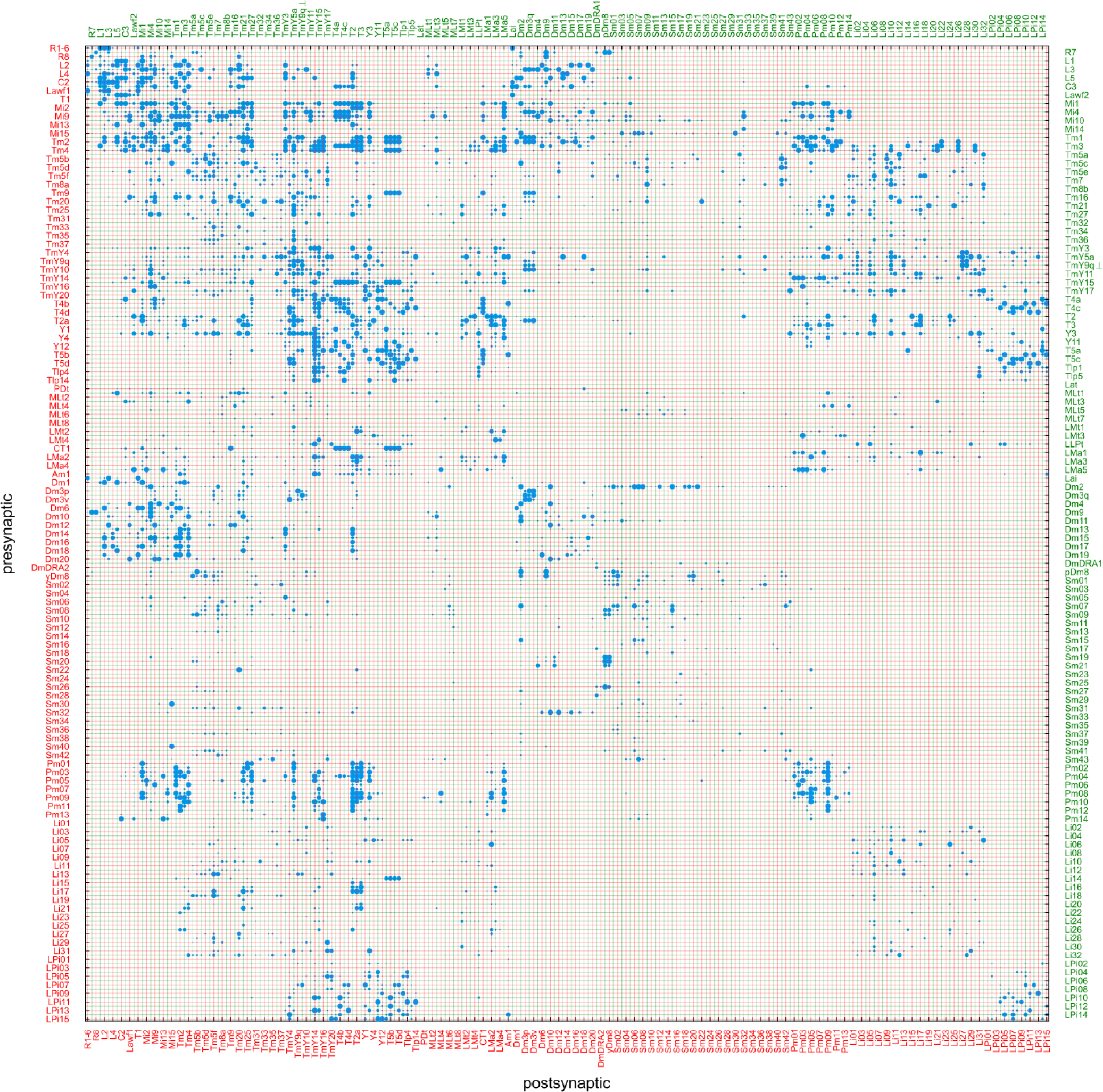
Type-to-type connectivity as a matrix. The number of synapses from one cell type to another is indicated by the area of the corresponding dot. Dot area saturates above 3600 synapses, to make weaker connections visible. For legibility, the type names alternate between left and right edges, and bottom and top edges, and are color coded to match the lines that are guides to the eye.

**Figure S5.**
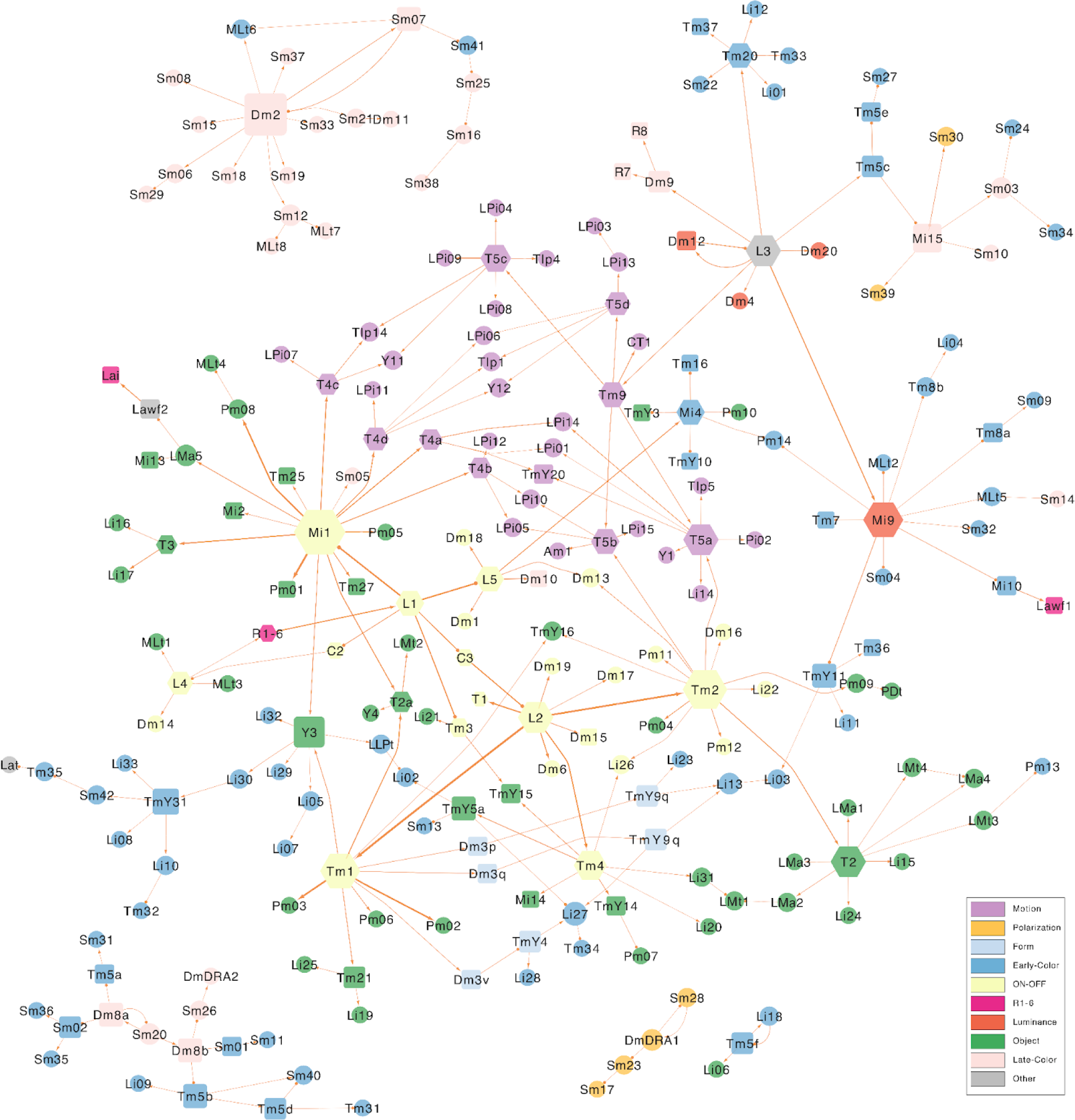
Wiring diagram of cell types (top input connections) Wiring diagram depicting top inputs for all cell types intrinsic to the optic lobe, as well as photoreceptors. Node size encodes the number of drawn connections, highlighting “hub” inputs. Node color indicates membership in the subsystems defined in the text (see legend). Node shape indicates number of cells (hexagons 800+, rectangles 100-799 and circles 1-99). Arrow tips indicate excitation and circle tips indicate inhibition.

**Figure S6.**
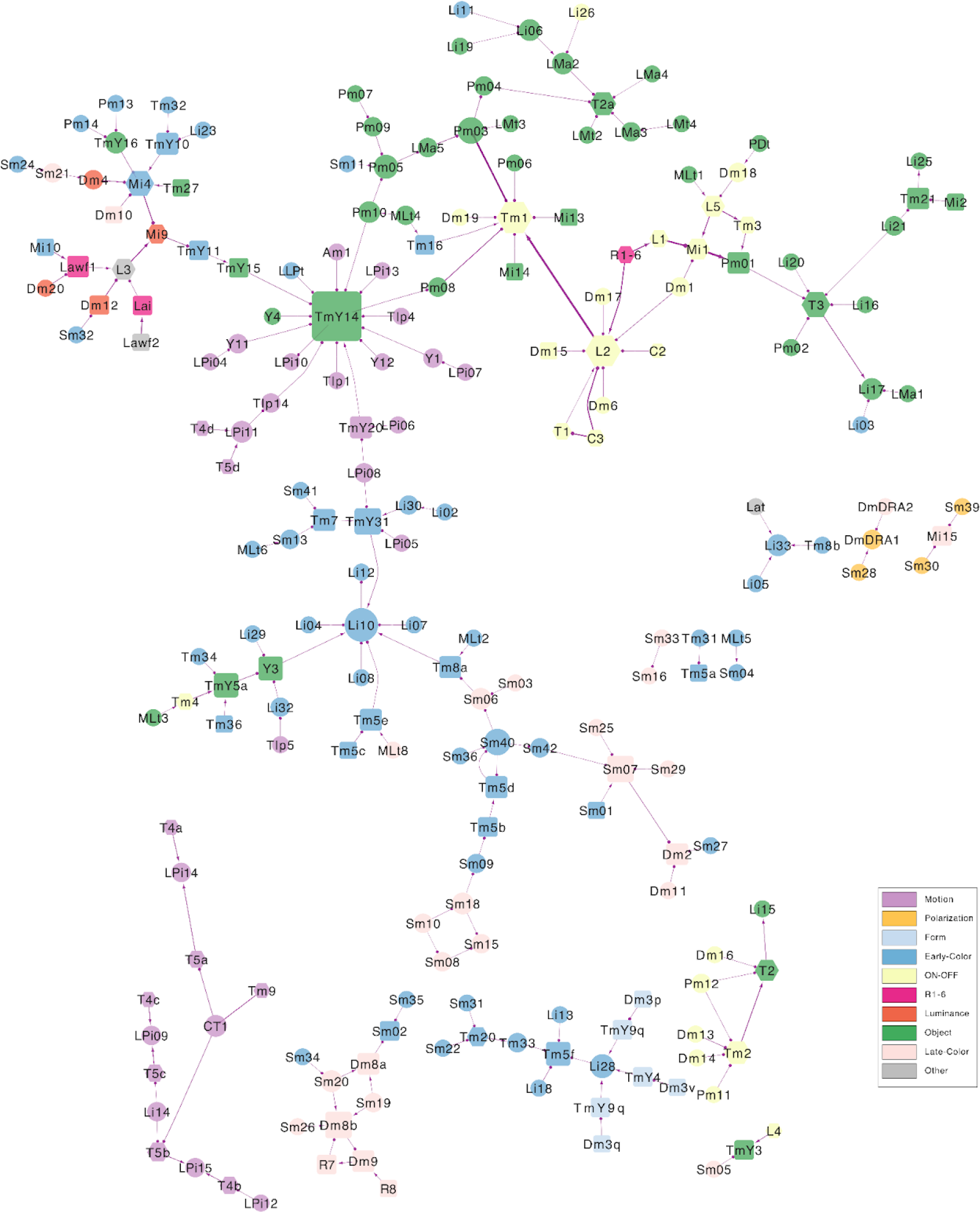
Wiring diagram of cell types (top output connections) Wiring diagram depicting top outputs. Node size encodes the number of drawn connections, highlighting “hub” outputs. Node color indicates membership in the subsystems defined in the text (see legend). Node shape indicates number of cells (hexagons 800+, rectangles 100-799 and circles 1-99). Arrow tips indicate excitation and circle tips indicate inhibition.

**Figure S7.**
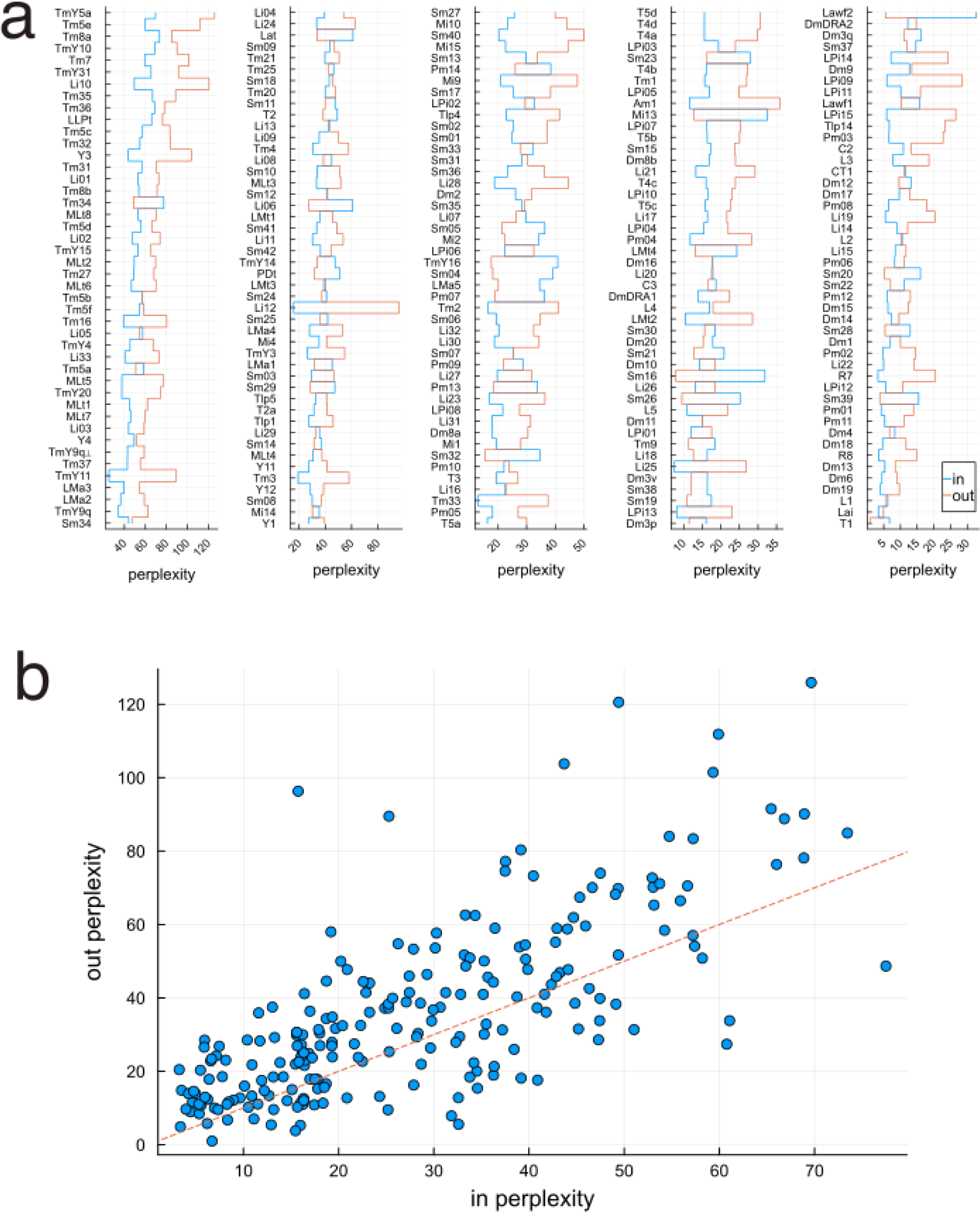
Input and output perplexity. (a) Input (blue) and output (red) perplexities. Types are ordered by the product of input and output perplexities. (b) Output and input perplexity are correlated. Output perplexity tends to exceed input perplexity (more points above red line).

**Figure S8.**
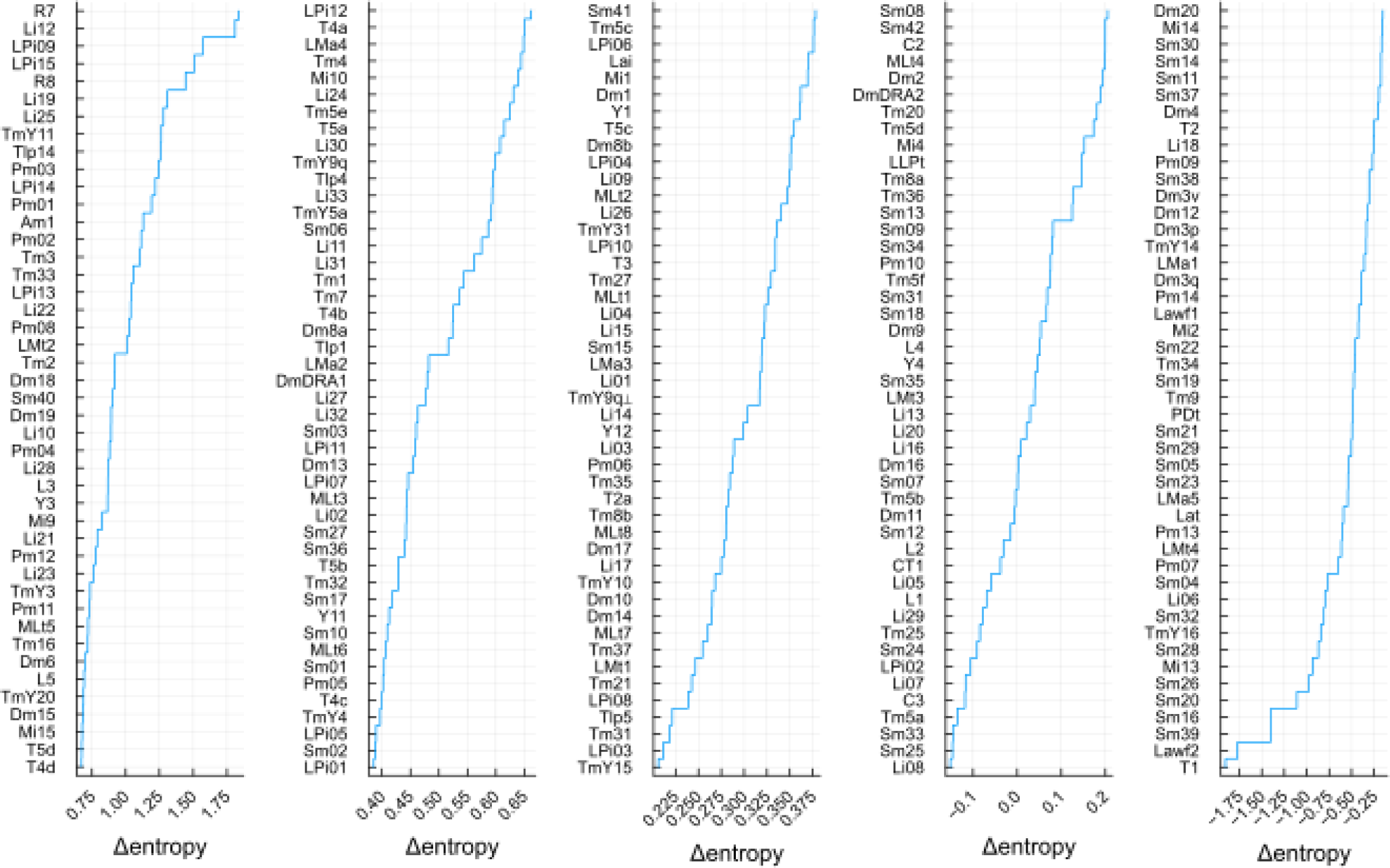
Difference between output and input entropies. The difference between output and input entropies (units of nats) quantifies the degree of divergence or convergence. This difference is equivalent to the logarithm of the ratio of out- and in-perplexities. The connectivity of the top types (top left) is more divergent, as the output entropy is greater than the input entropy. The connectivity of the bottom types (bottom right) is more convergent, as the input entropy is greater than the output entropy.

**Figure S9.**
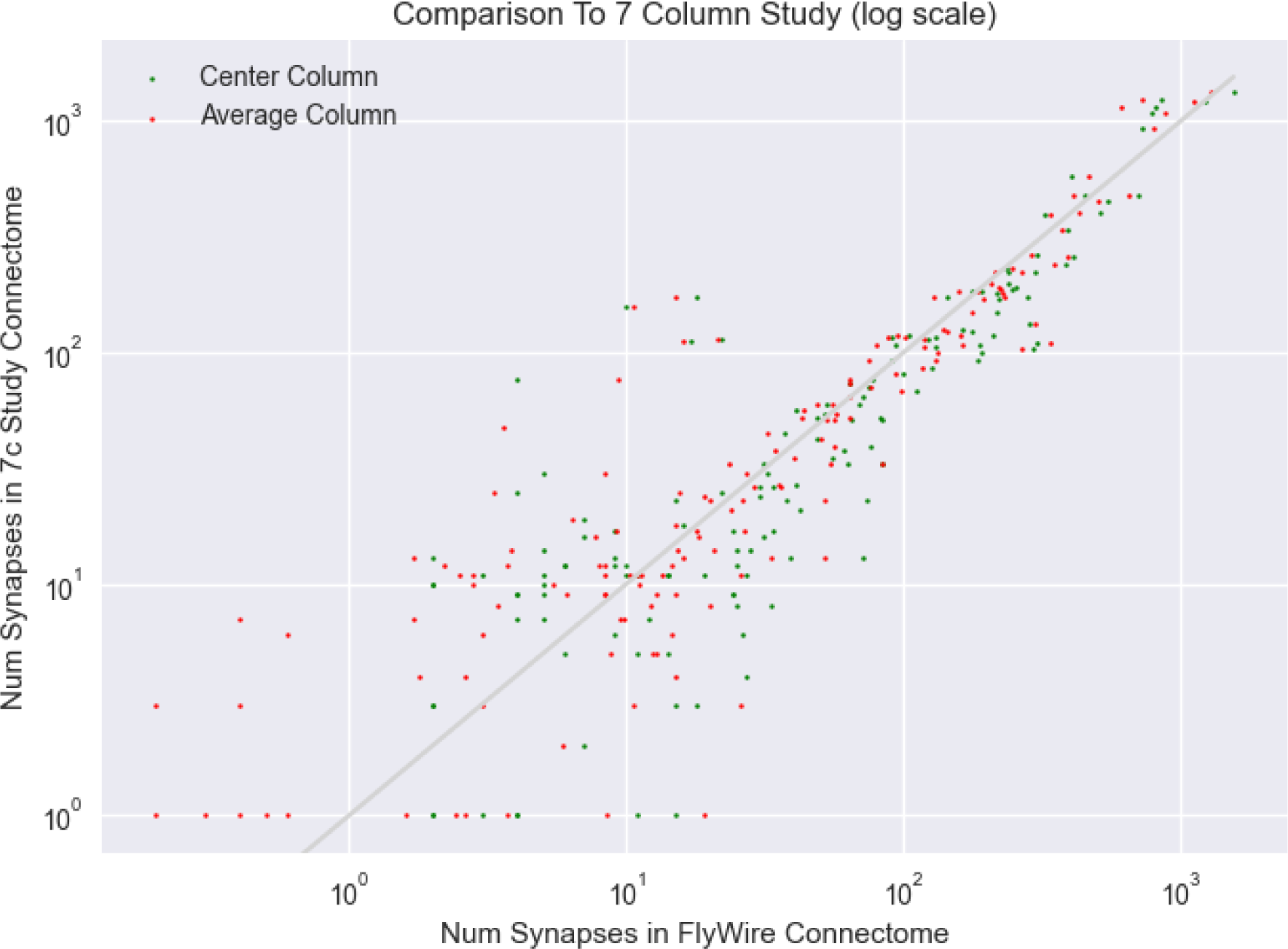
Comparison with seven-column reconstruction. We compared the synapse counts between type pairs to the corresponding synapse counts in the seven-column reconstruction. For this comparison we used the center column and its surrounding 6 columns from our dataset (green dots) as well as the average of 100 columns and their surrounding ones (red dots). Each point represents an ordered pair of types, and the number of synapses between them in the FlyWire connectome (X) and the seven-column reconstruction (Y). Correlation coefficients are 0.952 for the center + 6 columns and 0.954 for the average.

## Supplementary Tables

Table S1 (link) [Master table of cell types](https://drive.google.com/file/d/1xcefWmR5rVHjhlnLSt-LLYOG7i1eQUim/view?usp=drive_link)

**Table S2.** Type families and their properties.

|  | AFFINITY | LINKAGE | TYPES | CELLS | TRANS | NEUROPILS |
| --- | --- | --- | --- | --- | --- | --- |
| Centrifugal | Cross Neuropil | Axon Bearing / Columnar | 2 | 1511 | GABA | ME → ME, LA |
| Distal Medulla | Neuropil Intrinsic | Non-columnar | 21 | 3275 | GLUT | ME → ME |
| Distal Medulla Dorsal Rim Area | Neuropil Intrinsic | Non-columnar | 2 | 40 | GLUT | ME → ME |
| Lamina Intrinsic | Neuropil Intrinsic | Non-columnar | 1 | 231 |  | LA → LA |
| Lamina Monopolar | Cross Neuropil | Axon Bearing / Columnar | 5 | 3829 | ACH | ME, LA → ME |
| Lamina Tangential | Neuropil Intrinsic | Axon Bearing / Tangential | 1 | 12 |  | ME, LO, PLP, AME → LO |
| Lamina Wide Field | Cross Neuropil | Axon Bearing / Columnar | 2 | 318 | ACH | ME → LA |
| Lobula Intrinsic | Neuropil Intrinsic | Non-columnar | 33 | 759 | GABA | LO → LO |
| Lobula Lobula Plate Tangential | Cross Neuropil | Axon Bearing / Tangential | 1 | 36 | GABA | LO, LOP → LO, LOP |
| Lobula Medulla Amacrine | Cross Neuropil | Amacrine | 6 | 119 | GLUT | ME, LO → ME, LO |
| Lobula Medulla Tangential | Cross Neuropil | Axon Bearing / Tangential | 4 | 85 | GLUT | LO, ME → LO, ME |
| Lobula Plate Intrinsic | Neuropil Intrinsic | Non-columnar | 15 | 363 | GLUT | LOP → LOP |
| Medulla Intrinsic | Neuropil Intrinsic | Axon Bearing / Columnar | 8 | 3920 | ACH | ME → ME |
| Medulla Lobula Lobula Plate Amacrine | Cross Neuropil | Amacrine | 1 | 1 | GABA | LOP, ME, LO → LOP, LO, ME |
| Medulla Lobula Tangential | Cross Neuropil | Axon Bearing / Tangential | 8 | 295 | ACH | ME, LO → ME, LO |
| Photo Receptors | Neuropil Intrinsic | Axon Bearing / Columnar | 3 | 4727 |  | ME, LA → LA, ME |
| Proximal Distal Medulla Tangential | Neuropil Intrinsic | Axon Bearing / Tangential | 1 | 6 | DA | ME → ME |
| Proximal Medulla | Neuropil Intrinsic | Non-columnar | 14 | 597 | GABA | ME → ME |
| Serpentine Medulla | Neuropil Intrinsic | Non-columnar | 42 | 1479 | GABA | ME → ME |
| T Neuron | Cross Neuropil | Axon Bearing / Columnar | 12 | 9243 | ACH | ME, LO, LOP → LOP, LO, ME |
| Translobula Plate | Cross Neuropil | Axon Bearing / Columnar | 4 | 170 | GLUT | LOP → LOP, LO |
| Transmedullary | Cross Neuropil | Axon Bearing / Columnar | 26 | 8804 | ACH | ME, LO → ME, LO |
| Transmedullary Y | Cross Neuropil | Axon Bearing / Columnar | 12 | 2608 | ACH | ME, LOP, LO → LO, ME, LOP |
| Y Neuron | Cross Neuropil | Axon Bearing / Columnar | 5 | 630 | GLUT | LOP, ME, LO → LO, LOP, ME |

**Table S3.** Distribution of synapses over neuropils for each type family.

|  | ↑ LA | ↑ ME | ↑ LO | ↑ LOP | ↓ LA | ↓ ME | ↓ LO | ↓ LOP |
| --- | --- | --- | --- | --- | --- | --- | --- | --- |
| Centrifugal | 2,361 | 110,557 | 0 | 5 | 25,499 | 192,287 | 0 | 6 |
| Distal Medulla | 109 | 492,601 | 29 | 0 | 3 | 460,578 | 66 | 0 |
| Distal Medulla Dorsal Rim Area | 0 | 5,806 | 0 | 0 | 0 | 6,379 | 0 | 0 |
| Lamina Intrinsic | 53,890 | 14 | 0 | 0 | 4,729 | 0 | 0 | 0 |
| Lamina Monopolar | 118,561 | 335,433 | 0 | 0 | 4,639 | 1,017,745 | 0 | 0 |
| Lamina Tangential | 0 | 180 | 153 | 0 | 0 | 0 | 11 | 0 |
| Lamina Wide Field | 212 | 76,252 | 0 | 0 | 39,686 | 568 | 0 | 0 |
| Lobula Intrinsic | 0 | 165 | 465,268 | 242 | 0 | 5 | 311,293 | 25 |
| Lobula Lobula Plate Tangential | 0 | 0 | 18,515 | 1,542 | 0 | 0 | 13,572 | 6,752 |
| Lobula Medulla Amacrine | 0 | 214,538 | 108,008 | 1,179 | 0 | 135,892 | 111,694 | 116 |
| Lobula Medulla Tangential | 0 | 6,134 | 41,754 | 72 | 0 | 21,704 | 25,521 | 25 |
| Lobula Plate Intrinsic | 0 | 1 | 3 | 320,696 | 0 | 36 | 13 | 222,563 |
| Medulla Intrinsic | 318 | 529,814 | 2 | 0 | 1 | 1,075,708 | 16 | 0 |
| Medulla Lobula Lobula Plate Amacrine | 0 | 3,201 | 2,183 | 27,387 | 0 | 3,449 | 3,762 | 12,089 |
| Medulla Lobula Tangential | 0 | 33,069 | 2,147 | 10 | 0 | 28,264 | 10,922 | 1 |
| Photo Receptors | 1,335 | 21,213 | 0 | 0 | 108,479 | 26,500 | 0 | 0 |
| Proximal Distal Medulla Tangential | 0 | 2,978 | 0 | 0 | 0 | 1,186 | 0 | 0 |
| Proximal Medulla | 0 | 965,527 | 3 | 350 | 0 | 444,218 | 1 | 265 |
| Serpentine Medulla | 0 | 240,211 | 10 | 0 | 0 | 238,912 | 78 | 0 |
| T Neuron | 6,491 | 635,478 | 238,420 | 49,089 | 127 | 99,587 | 280,610 | 498,059 |
| Translobula Plate | 0 | 46 | 2,736 | 90,855 | 0 | 150 | 15,944 | 51,089 |
| Transmedullary | 0 | 936,228 | 91,635 | 920 | 0 | 1,024,221 | 579,930 | 1,667 |
| Transmedullary Y | 0 | 370,762 | 121,494 | 152,109 | 0 | 150,694 | 206,702 | 65,656 |
| Y Neuron | 0 | 82,196 | 23,720 | 84,681 | 0 | 55,853 | 78,629 | 56,527 |

**Table S4.**
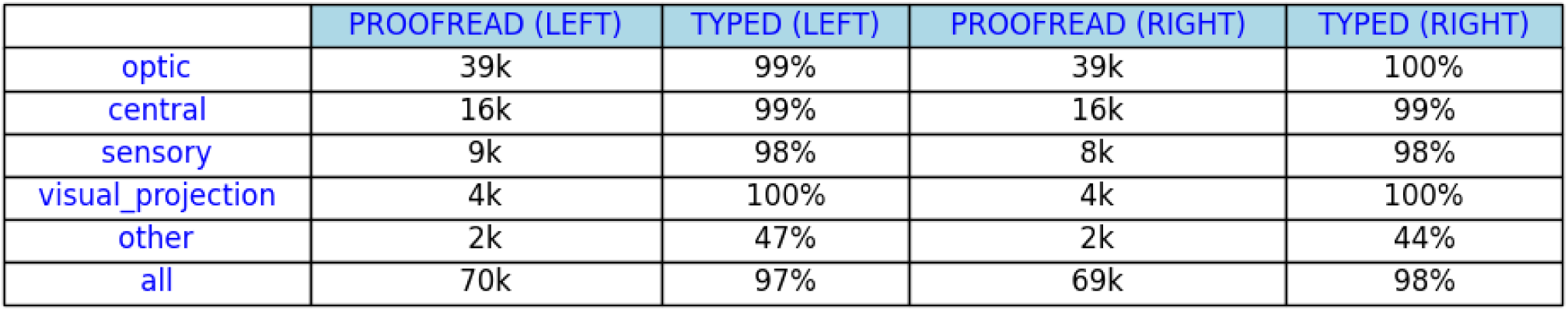
Status of cell typing. FlyWire connectome dataset as of October 2023.

|  | PROOFREAD (LEFT) | TYPED (LEFT) | PROOFREAD (RIGHT) | TYPED (RIGHT) |
| --- | --- | --- | --- | --- |
| optic | 39k | 99% | 39k | 100% |
| central | 16k | 99% | 16k | 99% |
| sensory | 9k | 98% | 8k | 98% |
| visual_projection | 4k | 100% | 4k | 100% |
| other | 2k | 47% | 2k | 44% |
| all | 70k | 97% | 69k | 98% |

Table S5 (link) [FlyWire Consortium (optic lobe)](https://docs.google.com/document/d/1Ag0s41C8--X8JCKmMk8BJhnj9Y1xYKIPIbcixJUrnfQ/edit?usp=drive_link)

## Supplementary Data

Data S1 (link) [Cards for all cell types](https://drive.google.com/drive/folders/1cWjwuDDW3yf6th3UQIgu6HZq0PJkDYtn?usp=drive_link)

Each card has 6 panels (clockwise): Logical connectivity predicate diagram, numeric stats, illustration of a typical cell, followed by 3D rendering of all cells from 3 orthogonal viewpoints.

Also available on [Codex](https://codex.flywire.ai/app/optic_lobe_catalog_static).

Data S2 (link) [Discriminating logical predicates for all types](https://drive.google.com/drive/folders/1E3h1GCl7Qjmhyfz0XLq-SxEZEAQW7aMi?usp=drive_link)

Each figure contains types from the same family (middle layer) with shared input attributes (left layer) and output attributes (right layer) that are sufficient for discriminating all types in the middle layer. Families with many types are split into multiple figures for clarity of presentation.

Data S3 (link) [Discriminating 2D projections for neuropil-intrinsic types](https://drive.google.com/drive/folders/15IqrAu-YT_u6emO_3kJUupn6DJ4Ge7_N?usp=drive_link)

Data S4 (link) [Input and output fractions for cell types](https://drive.google.com/drive/folders/1D6MsjSknyaWuAYwofZuG4QKKtyjMdXFo?usp=drive_link)

## Methods

### Reconstruction accuracy and completeness

The overall quality of our *Drosophila* brain reconstruction has been evaluated elsewhere (Dorkenwald et al. 2022; Dorkenwald, Matsliah, et al. 2023). Here we describe a few additional checks that are specific to the optic lobe. A small percentage of cells have eluded proofreading efforts. The worst cases are a few types with visible “bald spots” in a particular region with poor image quality, but underrrecovery is hardly visible for most types (Data S1). For a quantitative estimate of underrecovery, we can rely on the numerous types (see main text for definition). Each of these types is expected to contain roughly 800 cells, and all turn out to contain at least 720 cells in our reconstruction (Fig. 1d). This suggests that underrecovery is 10% at most, and typically less than that. The inner photoreceptors R7 and R8 are about 650 cells each, and the outer photoreceptors R1-6 total about 3400 in v783. However, more have been proofread and annotated in the latest version where we have a close number of eight photoreceptors per cartridge. Underrecovery is particularly severe in the numerous types located in the mid posterior side of the right optic lobe. In this region, we observed a notable narrowing and discontinuation of neuronal tracks. Many of these tracks appear to terminate within glial cells, suggesting a potential engulfment of neurons by glia. Additionally, the use of 40nm sections might be thicker to adequately capture these fine cellular details, possibly contributing to the observed underrecovery.

The connectivity between the numerous types can be compared with a previous reconstruction of seven medulla columns (S.-Y. Takemura et al. 2015), and there is good agreement (Fig. S9). While the agreement is reassuring, a cautionary note is that weaker connections in the type-type connectivity matrix (Fig. S4) could be artifactual, due to false positives of automated synapse detection. There are some heuristics for guessing whether a connection is artifactual, short of manually inspecting the original EM images. For example, one might distrust weak connections between cells, i.e., those with less than some threshold number of synapses. The choice of the threshold value depends on the context (Scheffer et al. 2020). For example, the flagship paper (Dorkenwald, Matsliah, et al. 2023) discarded connections with less than five synapses, a convention followed by the FlyWire Codex. The predicates of the present work apply a threshold of two synapses rather than five. The different thresholds were chosen because the central brain and optic lobes are very different contexts, as we now explain.

In the central brain, most cell types have cardinality 2 (cell and its mirror twin in the opposite hemisphere, Fig. S1e). In the hemibrain, the cardinality is typically reduced to one. Therefore if you would like to know whether there is a connection between cell type A and cell type B, you must decide based on only two or three examples of the ordered pair (A, B) in all the connectomic data that is so far available. Given the small sample size, it makes sense to set the threshold to a relatively high value.

In the optic lobe, on the other hand, there are often many examples of the ordered pair (A, B), because so many cell types have high cardinality. Therefore, if a connection is consistently found from type A to type B, one can have reasonable confidence even if the average number of synapses in the connection is not so high. That is why we set the threshold to a relatively low value in the optic lobe predicates. In particular, we have found that certain inhibitory types consistently make connections that involve relatively few synapses, and these connections seem real.

Another heuristic is to look for extreme asymmetry in the matrix. If the number of synapses from A to B is much larger than from B to A, the latter connection might be spurious. The reason is that the strong connection from A to B means the contact area between A and B is large, which means more opportunity for false positive synapses from B to A.

Finally, it may be known from other studies that a connection does not exist. For example, T1 cells lack output synapses (Meinertzhagen and O’Neil 1991; S.-Y. Takemura, Lu, and Meinertzhagen 2008). Therefore in our analyses, we typically regarded the few outgoing T1 synapses in our data as false positives and discarded them.

### Morphological cell typing

Our connectomic cell approach to typing is initially seeded with some set of types, in order to define the feature vectors for cells (Fig. 2a), after which the types are refined by computational methods. For the initial seeding, we relied on the time-honored approach of morphological cell typing, sometimes assisted by computational tools that analyzed connectivity. It is worth noting that “morphology” is a misnomer, because it refers to shape only, strictly speaking. Orientation and position are actually more fundamental properties because of their influence on stratification in neuropil layers. Therefore “single-cell anatomy” would be more accurate than “morphology,” though the latter is the standard term.

#### Stage 1: Crowdsourced annotation of known types

Annotations of optic lobe neurons were initially crowdsourced. The first annotators were volunteers from *Drosophila* labs. They were later joined by citizen scientists. At this stage, the annotation effort was mainly devoted to labeling cells of known types, especially the most numerous types.

##### *Drosophila* lab annotators

Labs that researched the *Drosophila* anterior visual pathway reconstructed and annotated medulla neurons that were upstream of the pathway. These included many of the medulla and lamina neurons discussed in this study. The annotated neurons were primarily Dm2, Mi15, R7, and R8, but also comprised various L, Dm, Mi, Tm, C, and Sm cells. Previously known neuron types were identified primarily by morphology and partially by connectivity. Annotators additionally found all Mi1 neurons in both hemispheres in order to find every medulla column. These Mi1 neurons were used to create the map of medulla layers based on Mi1 stratification in (Fischbach and Dittrich 1989). This layer map would later be used by the citizen scientists to identify medulla cell types.

##### Citizen scientists

The top 100 players from Eyewire (Kim et al. 2014) had been invited to proofread in FlyWire (Dorkenwald, Matsliah, et al. 2023). After three months of proofreading in the right optic lobe, they were encouraged to also label neurons when they felt confident. Most citizen scientists did a mixture of annotation and proofreading. Sometimes they annotated cells after proofreading, and other times searched for cells of a particular type to proofread.

Citizen scientists were provided with a visual guide to optic lobe cells sourced from (Fischbach and Dittrich 1989) and (Davis et al. 2020). FlyWire made available a 3D mesh overlay indicating the four main optic lobe neuropils. Visual identification was primarily based on single-cell anatomy (neuropils, stratification, and morphology). Initially labeling of type families (ie. Dm, Tm, Mi, etc.) was encouraged, especially for novices. Annotation of specific types (ie. Dm3, Tm2, etc.) developed over time. Type names from (Fischbach and Dittrich 1989) were learned by the citizen scientists from the visual guide. Use of canonical names was further enforced by a software tool that enabled easy selection and submission of pre-formatted type names.

Additional community resources (discussion board/forum, blog, shared Google drive, chat, dedicated email, and Twitch livestream) fostered an environment for sharing ideas and information between community members (citizen scientists, community managers, and researchers). Community managers answered questions, provided resources such as the visual guide, shared updates, performed troubleshooting and general organization of community activity. Daily stats including number of annotations submitted per individual were shared on the discussion board/forum to provide project progress. Live interaction, demonstrations and communal problem solving occurred during weekly Twitch video livestreams led by a community manager. The environment created by these resources allowed citizen scientists to self-organize in several ways: community driven information sharing, programmatic tools, and “farms.”

##### Community-driven information sharing

Citizen scientists created a comprehensive guide with text and screenshots that expanded on the visual guide. They also found and studied any publicly available scientific literature or resources regarding the optic lobe. They shared findings on the [Discussion Board](https://discuss.flywire.ai/), which as of October 10, 2023 had over 2,500 posts. Community managers interacted with citizen scientists by sharing findings from the scientific literature, consulting *Drosophila* specialists on FlyWire, and providing feedback.

##### Programmatic Tools

Programmatic tools were created to help with searching for cells of the same type. One important script traced partners-of-partners, i.e., source cell → downstream partners → their upstream partners, or source cell → upstream partners → their downstream partners. This was based on the assumption that cells of the same type will probably synapse with the same target cells, which often turned out to be true. The tool could either look for partners-of-all-partners or partners-of-any-partners. The resulting lists of cells could be very long, and were filtered by excluding cells that had already been identified, or excluding segments with small sizes or low ID numbers (which had likely not yet been proofread).

Another tool created from LPTCs (e.g. HS, VS, H1) aided definition of layers in the lobula plate. This facilitated identification of various cell types, especially T4 and T5.

##### Cell “Farms”

Citizen scientists created “farms” in FlyWire or Neuroglancer with all the found cells of a given type visible. Farms showed visually where cells still remained to be found. If they found a bald spot, a popular method to find missing cells was to move the 2D plane in that place and add segments to the farm one after another in search of cells of the correct type. Farms also helped with identifying cells near the edges of neuropils, where neurons are usually deformed. Having a view of all other cells of the same type made it possible to extrapolate to how a cell at the edge should look.

#### Stage 2: Centralized annotation and discovery of new types

A team of image analysts at Princeton finished the annotation of the remaining cells in known types, and also discovered new types. We began by utilizing community annotations from each type. To ensure accuracy, these annotations were initially cross-referenced with existing literature to confirm their validity. Once validated, these cells were used to query various Codex search tools that returned previously unannotated cells exhibiting connectivity similar to that of the cell in the query. The hits from the search query were evaluated by morphology and stratification to confirm match with the target cell type. In some cases where cell type distinctions were uncertain, predicted neurotransmitters (Eckstein et al. 2023) were used for additional guidance. This process allowed us to create a preliminary clustering of all previously known and new types.

### Connectomic cell typing

Eventually morphology became insufficient for further progress. Expert annotators, for example, struggled to classify Tm5 cells into the three known types, not knowing that there would turn out to be six Tm5 types. At this point, we were forced to transition to connectomic cell typing. In retrospect, this transition could have been made much earlier. As mentioned above, connectomic cell typing must be seeded with an initial set of types, but the seeding did not have to be as thorough as it ended up. We leave for future work the challenge of extending the connectomic approach so it can be used from start to finish.

#### Stage 3: Connectivity-based splitting and merging of types and auto-correction

We used computational methods to split types that could not be properly split in Stage 2. Some candidates for splitting (e.g. Tm5) were suggested by the image analysts of Stage 2. Some candidates were suspicious because they contained so many cells. Finally, some candidates were scrutinized because their type radii were large. We applied hierarchical clustering with average linkage, and accepted the splits if they did not violate the tiling principle as described in Spatial Coverage.

We also applied computational methods to merge types that had been improperly split in Stage 2. Here the candidates were types with low spatial coverage of the visual field, or types that were suspiciously close in the dendrogram of cell types (Fig. 2c). Merge decisions were made by hierarchical clustering of cells from types that were candidates for merging, and validated if they improved spatial coverage.

Once we arrived at the final list of types, we estimated the “center” of each type using the element-wise trimmed mean. Then for every cell we computed the nearest type center by Jaccard distance. For 98% of the cells, the nearest type center coincided with the assigned type. We sampled some disagreements and reviewed them manually. In the majority of cases, the algorithm was correct, and the human annotators had made errors, usually of inattention. The remaining cases were mostly attributable to proofreading errors. There were also cases in which type centers had been contaminated by human-misassigned cells (see Morphological Variation), which in turn led to more misassignment by the algorithm. After addressing these issues, we applied the automatic corrections to all but 0.1% of cells, which were rejected using distance thresholds.

### Validation

Based on the auto-correction procedure, we estimate that our cell type assignments are between 98% and 99.9% accurate. For another measure of the quality of our cell typing, we computed the “radius” of each type, defined as the average distance from its cells to its center. Here we computed the center by approximately minimizing the sum of Jaccard distances from each cell in the type to the center (Computational concepts). A large type radius can be a sign that the type contains dissimilar cells, and should be split. For our final types, the radii vary, but almost all lie below 0.6 (Fig. S3a). Lat is one exception, and may possibly deserve to be split (see Cross-neuropil tangential and amacrine). The type radii are essentially the same, whether or not boundary types are included in the feature vector (data not shown).

#### Discrimination with logical predicates

Because the feature vector is rather high dimensional, it would be helpful to have simpler insights into what makes a type. One approach is to find a set of simple logical predicates based on connectivity that predict type membership with high accuracy. For a given cell, we define the attribute “is connected to input type *t*” as meaning that the cell receives at least one connection from some cell of type *t*. Similarly, the attribute “is connected to output type *t*” means that the cell makes at least one connection onto some cell of type *t*.

An optimal predicate is constructed for each type that consists of 2 tuples: input types and output types. Both tuples are limited to size 5 at most, and they are optimal with respect to the F-score of their prediction of the subject type, defined as follows:

- Recall of a predicate for type *T* is the ratio of true positive predictions (cells matching the predicate) to the total number of true positives (cells of type *T*). It measures the predicate’s ability to identify all positive instances of a given type.
- Precision is the ratio of true positive predictions (predictions that are indeed of type T) to the total number of positive predictions made by the logical predicate.
- F-score is the harmonic mean of precision and recall - a single metric that combines both precision and recall into one value.

On a high level, the process for computing the predicates is exhaustive - for each type we look for all possible combinations of input type tuples and output type tuples and compute their precision, recall and f-score. Few optimization techniques are used to speed up this computation, by calculating minimum precision and recall thresholds from the current best candidate predicate and pruning many tuples early.

For example, the logical predicate “is connected to input type Tm9 and output type Am1 and output type LPi15” predicts T5b cells with 99% precision and 99% recall. For all but three of the identified types we found a logical predicate with 5 or fewer input / output attributes that predicts type membership with average f-score 0.93, weighted by the number of cells in type (see Figure S4 and Table S1). Some of the attributes in a predicate are the top most connected partner types, but this is not necessarily the case. The attributes are distinctive partners, which are not always the most connected partners. The predicate for each type is shown on its card in Data S1. For each family, the predicates for all types can be shown together in a single graph containing all relevant attributes (Data S2).

We experimented with searching for predicates after randomly shuffling a small fraction of types (namely, swapping types for 5% of randomly picked pairs of neurons). We found that precision and recall of the best predicates dropped dramatically, suggesting that we are not overfitting. This was expected because the predicates are short.

We also measured the drop in the quality of predicates if excluding boundary types (where the predicates are allowed to contain intrinsic types only). As is the case with the clustering metrics, the impact on predicates is marginal (weighted mean F-score drops from 0.93 to 0.92; Methods).

#### Discrimination with two-dimensional projections

Another approach to interpretability is to look at low-dimensional projections of the 2*T*-dimensional feature vector. For each cell type, we select a small subset of dimensions that suffice to accurately discriminate that type from other types (Fig. S3c). Here we normalize the feature vector so that its elements represent the “fraction of input synapses received from type *t*” or “fraction of output synapses sent to type *t*.” In these normalized quantities, the denominator is the total number of all input or output synapses, not just the synapses with other neurons intrinsic to the optic lobe.

For example, we can visualize all cells in the Pm family in the two-dimensional space of C3 input fraction and TmY3 output fraction (Fig. S3c). In this space, Pm04 (Pm⇐C3⇒TmY3) cells are well-separated from other Pm cells, and can be discriminated with 100% accuracy by “C3 input fraction greater than 0.01 and TmY3 output fraction greater than 0.01.”

More generally, a cell type discriminator is based on thresholding a set of input and output fractions, and taking the conjunction of the result. The search for a discriminator finds a set of dimensions, along with threshold values for the dimensions. To simplify the search, we require that the cell type be discriminated only from other types in the same neuropil family, rather than from all other types. Under these conditions, it almost always suffices to use just two dimensions of the normalized feature vector.

This approach was applied to interneuron families, which contain many types that can be difficult to distinguish. Discriminating 2D projections for each interneuron type are provided in Data S3. Many though not all discriminations are highly accurate. Both intrinsic and boundary types are included as discriminative features.

### Computational concepts

#### Connectivity: cell-to-cell, type-to-cell, cell-to-type, and type-to-type

Connectomic cell typing is based on the feature vectors of Fig. 2a. The feature vectors are defined formally as follows. Define a (weighted) cell-to-cell connectivity matrix *w_ij_*, as the number of synapses from neuron *i* to neuron *j*. The synapse out-degree and in-degree of neuron *i* are:

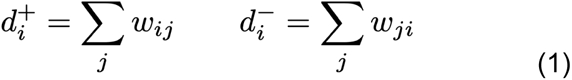

The sums are over all neurons in the brain. If neuron *i* is a cell intrinsic to one optic lobe, the only nonvanishing terms in the sums are due to the intrinsic and boundary neurons for that optic lobe.

Let *A_it_* be the 0-1 matrix that assigns neuron *i* to type *t*. The column and row sums of the assignment matrix satisfy

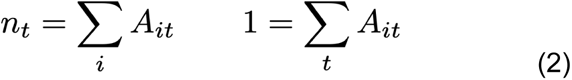

where *n_t_* is the number of cells assigned to type *t*. The assignment matrix may include only intrinsic types (Fig. 2a), or both intrinsic and boundary types.

The cell-to-type connectivity matrix *O_it_* is the number of output synapses from neuron *i* to neurons of type *t*,

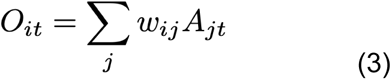

and the type-to-cell connectivity matrix *I_tj_* is the number of input synapses from neurons of type *t* onto neuron *j*,

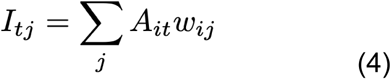

The *i*th row and ith column of these matrices are concatenated to form the feature vector (Fig. 2a) used for computing Jaccard similarities and distances. These matrices are normalized by degree to yield the output and input fractions of cell 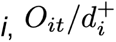 and 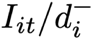. Elements of these matrices are used for the discriminating 2D projections (Fig. S3c).

The type-to-type connectivity matrix is the number of synapses from neurons of type *s* to neurons of type *t*,

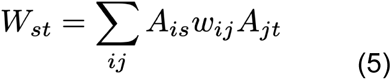

The synapse degree of type *t* is the sum of the degrees of the cells in type *t*,

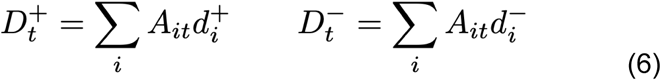

Normalizing by degree yields the output fractions of type 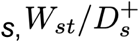, where *t* runs from 1 to *T*. The input fractions of type *t* are similarly given by 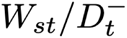, where *s* runs from 1 to *T*. Selected output and input fractions of types are shown in Data S4.

Alternatively, the feature vectors can be based on connection number rather than synapse number, where a connection is defined as two or more synapses from one neuron to another. Such a threshold is intended to suppress noise due to false positives in the automated synapse detection. Synapse number and connection number give similar results, and we use both in our analyses.

The cell type feature vector is obtained by normalizing and then concatenating the corresponding row and column of the type-to-type connectivity matrix.

#### Similarity and distance measures

The weighted Jaccard similarity between feature vectors **x** and **y** is defined by

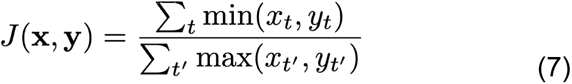

and the weighted Jaccard distance *d*(**x**, **y**) is defined as one minus the weighted Jaccard similarity. These quantities are bounded between zero and one since our feature vectors are nonnegative. In our cell typing efforts, we have found empirically that Jaccard similarity works better than cosine similarity when feature vectors are sparse.

#### Cluster centers

Given a set of feature vectors **x**^a^, the center **c** can be defined as the vector minimizing

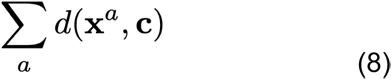

This cost function is convex, since *d* is a metric satisfying the triangle inequality. Therefore the cost function has a unique minimum. We used various approximate methods to minimize the cost function.

For auto-correction of type assignments, we used the element-wise trimmed mean. We found empirically that this gave good robustness to noise from false synapse detections. For the type radii, we used a coordinate descent approach, minimizing the cost function with respect to each *c_i_* in turn. The loop included every *i* for which some *x_i_* was nonzero. This converged within a few iterations of the loop.

### Axon versus dendrite

The main text (Class, Family, and Type) used the term “axon.” An axon is defined as some portion of the neuron with a high ratio of presynapses to postsynapses. This ratio might be high in an absolute sense. Or the ratio in the axon might only be high relative to the ratio elsewhere in the neuron (the dendrite). In either case, the axon is typically not a pure output element, but has some postsynapses as well as presynapses. For many types it is obvious whether there is an axon, but for a few types we have made judgment calls. Even without examining synapses, the axon can often be recognized from the presence of varicosities, which are presynaptic boutons.

### Columnar neurons

(Fischbach and Dittrich 1989) defined 13 columnar families (Fig. 1a). Those consisting exclusively of “numerous” (∼800 cells) types include L (lamina to medulla), C (medulla to lamina), T1 (distal medulla to lamina), T2 (distal and proximal medulla to lobula), T3 (proximal medulla to lobula), T4 (proximal medulla to lobula plate), and T5 (proximal medulla to lobula and lobula plate). We follow the convention of grouping the less numerous Lawf1 (distal medulla to lamina) and Lawf2 (proximal and distal medulla to lamina) types in the same family. Although T1 shares the same neuropils with Lawf1, T1 lacks output synapses (Meinertzhagen and O’Neil 1991; S.-Y. Takemura, Lu, and Meinertzhagen 2008), so it is an outlier and deserves to be a separate family. We regard distal and proximal medulla as two separate neuropils following (Fischbach and Dittrich 1989).

#### Mi Medulla intrinsic

(Fischbach and Dittrich 1989) defined Mi as projecting from distal to proximal medulla. Mi contains both numerous and less numerous types. We identified five (Mi1, 2, 4, 9, 10) of the dozen Mi types defined by (Fischbach and Dittrich 1989), and three (Mi13, 14, 15) types uncovered by EM reconstruction (S.-Y. Takemura et al. 2013). Mi1, Mi4, and Mi9 are consistent with the classical definition, but Mi13 projects from proximal to distal medulla. Other Mi types are less polarized, and narrow-field amacrine might be more accurate than columnar. Nevertheless we will adhere to the convention that they are columnar. Narrow-field amacrine cells are also found in the Sm family, and exist in the mammalian retina (Marc et al. 2014).

#### Tm Transmedullary

As classically defined (Fischbach and Dittrich 1989), Tm cells project from the distal medulla to the lobula. Tm1 through Tm26 and Tm28 were defined by (Fischbach and Dittrich 1989), and Tm27/Tm27Y was reported by (Hasegawa et al. 2011). We were able to identify Tm1, 2, 3, 4, 7, 9, 16, 20, 21, 25, and 27. We split Tm5 into six types, and Tm8 into two types. We merged Tm6 and Tm21 into a single type Tm21. We prefer the latter name because the cells more closely match the Tm21 stratification as drawn by (Fischbach and Dittrich 1989). Tm1a and Tm4a were defined as morphological variants (Fischbach and Dittrich 1989), but we have found that they do not differ in connectivity and are not common, so we have merged them into Tm1 and Tm4, respectively. We merged Tm27Y into Tm27 (Hasegawa et al. 2011), and Tm5Y and TmY5 into TmY5a (Fischbach and Dittrich 1989; Morante and Desplan 2008). These morphological distinctions originally arose because the projection into the lobula plate, the differentiator between Tm and TmY, can vary across cells in a type. We added Tm31 to Tm37, which project from the serpentine medulla to the lobula. We moved Tm23 and Tm24 to the Lobula intrinsic (Li) family. They were originally classified as Tm because their cell bodies are in the distal rind of the medulla, and they send a neurite along the columnar axis of the medulla to reach the lobula (Fischbach and Dittrich 1989). However, they do not form synapses in the medulla, so we regard them as Li neurons in spite of their soma locations. Overall, roughly half of the 26 types in the Tm family are novel.

#### TmY

Transmedullary Y (TmY) cells project from the distal medulla to the lobula and lobula plate. The “Y” refers to the divergence of branches to the lobula and lobula plate. TmY1 to TmY13 were defined by (Fischbach and Dittrich 1989), TmY5a by (Morante and Desplan 2008; Fischbach and Dittrich 1989), TmY14 by (S.-Y. Takemura et al. 2013), TmY15 by (Takemura et al. 2017), TmY16, TmY18, and TmY20 by (Shinomiya et al. 2019). We identified TmY3, TmY4, TmY5a, TmY10, TmY11, TmY14, TmY15, TmY16, and TmY20. We divided TmY9 into two types, as discussed in a companion paper (H. Sebastian Seung 2023). We added a new type TmY31.

#### Y

Y cells project from the proximal medulla to the lobula and lobula plate. They are similar to TmY cells, but the latter traverse both the distal and proximal medulla (Fischbach and Dittrich 1989). Y1 and Y3 to Y6 were defined by (Fischbach and Dittrich 1989). Y11 and Y12 were defined by (Shinomiya et al. 2022). We have identified Y1, Y3, Y4, Y11, and Y12 in our reconstruction, and have not found any new Y types. Y1, Y11, and Y12 have the majority of their synapses in the lobula plate, and are assigned to the motion subsystem. Y3 and Y4 have few synapses in the lobula plate, and are assigned to the object subsystem (Fig. 2). Y3 is more numerous (∼300 cells) than Y4, and is the only Y type that is predicted cholinergic.

#### Tlp Translobula plate

A translobula plate neuron projects from the lobula plate to the lobula. Tlp1 through Tlp5 were defined by (Fischbach and Dittrich 1989), and Tlp11 through Tlp14 by (Shinomiya et al. 2022). We have identified Tlp1, Tlp4, Tlp5, and Tlp14. We propose that the names Tlp11, Tlp12, and Tlp13 be retired (Shinomiya et al. 2022), as these types can now be unambiguously identified with Tlp5, Tlp1, and Tlp4 respectively.

### Interneurons

A local interneuron is defined as being completely confined to a single neuropil (Fig. 1b). Interneurons make up the majority of types, but a minority of cells (Fig. 1e). Lai is the only lamina interneuron. Dm and Pm interneurons (Fischbach and Dittrich 1989) stratify in the distal or proximal medulla, respectively. We have more than doubled the number of Pm types, and slightly increased the number of Dm types. We introduce the Serpentine medulla (Sm) family, which is almost completely new and contains more types than any other family (Fig. 1f). Li and LPi interneurons stratify in the lobula or lobula plate, respectively. Interneurons are usually amacrine and inhibitory (GABA or glutamate), but some are tangential or cholinergic. Interneurons are often wide field but some are narrow field.

#### Dm Distal Medulla

Dm1 through Dm8 were defined by (Fischbach and Dittrich 1989), Dm9 and 10 by (S.-Y. Takemura et al. 2013), and Dm11 through Dm20 by (Nern, Pfeiffer, and Rubin 2015). We do not observe Dm5 and Dm7, consistent with a previous study (Nern, Pfeiffer, and Rubin 2015). Most types are predicted to secrete glutamate or GABA, but there are also a few cholinergic types (Table S1). To Dm3p and Dm3q (Nern, Pfeiffer, and Rubin 2015; Özel et al. 2021; Kurmangaliyev et al. 2020), we added a third type Dm3v (Data S1). We split Dm8 into Dm8a and Dm8b (see Correspondences).

#### Dm Dorsal Rim Area

The Dorsal Rim Area (DRA) differs from the rest of the retina in its organization of inner photoreceptors. Photoreceptors in non-DRA and DRA differ in their axonal target layers and output cell types (von Philipsborn and Labhart 1990; Sancer et al. 2019). Specifically, DRA-R7 connects with DmDRA1 whereas DRA-R8 connects to DmDRA2 (Sancer et al. 2019; Hardcastle et al. 2021). These distinctive connectivity patterns result in DmDRA1 and DmDRA2 types exhibiting an arched coverage primarily in the M6 layer of the dorsal medulla (Fig. 9b). R7-DRA and R8-DRA are incompletely annotated at present, and this will be rectified in a future release. DmDRA1 receives R7 input, but sits squarely in M7. This could be regarded as an Sm type, but we have chosen not to change the name for historical reasons.

#### Pm Proximal medulla

Pm1, 1a, and 2 (Fischbach and Dittrich 1989) were each split into two types. Pm3 and 4 were defined by (Nern, Pfeiffer, and Rubin 2015). We additionally defined six new Pm types, for a total of 14 Pm types, numbered Pm01 to Pm14 in order of increasing average cell volume. The new names can be distinguished from the old ones by the presence of leading zeros. All are predicted GABAergic.

Pm1 was split into Pm06 (Pm⇒Mi9) and Pm04 (Pm⇐C3⇒TmY3). Pm1a was split into Pm02 (Pm⇐Tm1⇒T3) and Pm01 (Pm⇐Mi1⇒Tm25). Pm2 was split into Pm03 (Pm⇐Tm1⇒Tm2) and Pm08 (Pm⇒Mi1⇒Tm1).

#### Sm Serpentine medulla

According to the original scheme of (Fischbach and Dittrich 1989), Dm and Pm interneurons stratify on the distal or proximal side, respectively, of the serpentine layer (M7) of the medulla. Many interneuron types turn out to have significant stratification in the serpentine layer, and these borderline cases constitute a large new “Serpentine medulla” (Sm) family of interneurons, almost all new. They have been named Sm01 to Sm42, in order of increasing average cell volume. The Sm family includes types recently named “medulla tangential intrinsic” (Kind et al. 2021). We avoid using this term indiscriminately because some Sm types are tangential while others are amacrine. Some Sm types spill over from M7 into the distal or proximal medulla, and a few reach from M7 to more distant medulla layers.

Sm stratification in M7 has functional implications. First, Sm types are positioned to communicate with the medulla tangential (Mt) cells and other boundary types that are important conduits of information in and out of the optic lobe (Data S4). Second, Sm types are positioned to communicate with the inner photoreceptor terminals, which are in M6 or at the edge of M7. Consequently many Sm types are involved in the processing of chromatic stimuli, and end up being assigned to the color subsystem.

The Sm family more than doubles the number of medulla intrinsic types, relative to the old scheme with only Pm and Dm. The Sm family might be related to the M6-LN class of neuron defined by (Chin et al. 2014). The correspondence is unclear because M6-LN neurons are defined to stratify in M6, while Sm mainly stratifies in M7. But some Sm types stratify at the border between M6 and M7, and therefore could be compatible with the M6-LN description.

#### Li Lobula intrinsic

Two lobula intrinsic types (Li1 and Li2) were defined by (Fischbach and Dittrich 1989) and 12 more (Li11 through 20 and mALC1 and mALC2) by (Scheffer et al. 2020). Of these we have confirmed Li2, Li12, Li16, mALC1, and mALC2. Li11 has been split into two types (see Morphological Variation), and Li18 into three types. We identified 21 additional Li types, but have not been able to make conclusive correspondences with previously identified types. As mentioned earlier, we transfer Tm23 and Tm24 (Fischbach and Dittrich 1989) from the Tm to the Li family. This amounts to a total of 33 Li types, which have been named Li01 to Li33 in order of increasing average cell volume.

Collisions with the names of (Fischbach and Dittrich 1989) are avoided by the presence of leading zeros in our new names. Collisions with the hemibrain names Li11 to Li20 (Scheffer et al. 2020) do not worry us because these particular names have been used by few or no publications so far, so there is little cost associated with name changes. In any case, we were only able to establish conclusive correspondences for a minority of the hemibrain Li11 to Li20 types, which are detailed in Table S1.

We considered merging Li04 (Li⇐Tm8b⇒aMe20) and Li07 (Li⇐mALC5⇒Tm16), but their connectivity is quite different. Furthermore, in a hierarchical agglomerative clustering, Li07 (Li⇐mALC5⇒Tm16) merges with Li08 (Li⇐TmY17⇒TmY10) rather than Li04 (Li⇐Tm8b⇒aMe20).

#### LPi Lobula plate intrinsic

LPi1-2 and 2-1 were defined by (Shinomiya et al. 2022), generalizing previous definitions of LPi3-4 and 4-3 by (Mauss et al. 2015). Other types reported were LPi2b and LPi34-12 (Shinomiya et al. 2022). (We are not counting fragments for which correspondences are not easy to establish.) We have added nine new types, for a total of 15 LPi types.

LPi names were originally based on stratification in layers 1 through 4 of the lobula plate. Now that LPi types have multiplied, stratification is no longer sufficient for naming. The naming system could be salvaged by adding letters to distinguish between cells of different sizes. For example, LPi15 and LPi05 could be called LPi2-1f and LPi2-1s, where “f” means full-field and “s” means small. For simplicity and brevity, we instead chose the names LPi01 to LPi15, in order of increasing average cell volume. Correspondences with old stratification-based names are detailed in Table S1.

### Cross-neuropil tangential and amacrine

Most types that span multiple neuropils are columnar. One tangential type that spans multiple neuropils inside the optic lobe was previously described: Lat has a tangential axon that projects from the medulla to the lamina (Fischbach and Dittrich 1989). There is some heterogeneity in the Lat population, as reflected in the relatively large type radius (Fig. S3a). We have decided to leave splitting for future work, as Lat has many dense core vesicles that are presently unannotated.

Here we introduce two new families of cross-neuropil types that are tangential (MLt1-8 and LMt1-4), and one that is amacrine (LMa1-5). Along with two new tangential families (PDt, LLPt) that contain only single types, and the known CT1 and Am1 types, that is a total of 21 cross-neuropil types that are non-columnar (Fig. 1c). Each of the new types (except PDt with 6 cells) contains between 10 and 100 cells.

The tangential types connect neuropils within one optic lobe and do not leave the optic lobe. Our usage of the term “tangential” focuses on axonal orientation only. It should not be misunderstood to imply a wide-field neuron that projects out of the optic lobe, which is the case for the well-known lobula plate tangential cells (LPTCs) or lobula tangential cells (LTs). The term “tangential” presupposes that we can identify an axonal arbor for the cell (see Axon versus Dendrite).

#### PDt Proximal to Distal medulla tangential

We found one tangential type that projects from proximal to distal medulla (Data S1).

#### MLt Medulla-Lobula tangential

(Kind et al. 2021) described ML1, a tangential neuron projecting from medulla to lobula. We will refer to this type as MLt1, and have discovered more types of the same family, MLt2 through MLt8. Mlt1 and Mlt2 dendrites span both distal and proximal medulla, and Mlt3 dendrites are in the distal medulla, so MLt1 through MLt3 receive L input (Data S1, S4). Mlt4 dendrites are in the proximal medulla (Data S1). Mlt5 through Mlt8 have substantial arbor overlap with the serpentine layer M7 (Data S1), and are therefore connected with many Sm types to be discussed later on (Data S4). Interaction between MLt types is fairly weak, with the exception of MLt7 to MLt5 (Data S4). MLt7 and MLt8 are restricted to the dorsal and dorsal rim areas.

#### LMt Lobula-Medulla tangential

We identified four tangential types (LMt1 through LMt4) that project from lobula to medulla. Their axonal arbors are all in the proximal medulla (Data S1), thinly stratified near layer M7, so they have many Pm targets (Data S4). Only LMt4 exhibits partial coverage.

#### LLPt Lobula-Lobula Plate tangential

We discovered one tangential type that projected from lobula to lobula plate, and called it LLPt. This is just a single type, rather than a family.

#### LMa Lobula Medulla amacrine

We discovered four amacrine types that extend over lobula and medulla. LMa1 through LMa4 are coupled with T2, T2a, and T3, and LMa4 and LMa3 synapse onto T4 and T5 (Data S4). The LMa family could be said to include CT1, a known amacrine cell that also extends over both the lobula and medulla. The new LMa types, however, consist of smaller cells that each cover a fraction of the visual field, whereas CT1 is a wide-field cell.

#### MLLPa Medulla Lobula Lobula Plate amacrine

(Shinomiya et al. 2022) reported Am1, a wide-field amacrine cell that extends over the medulla, lobula, and lobula plate. We found no other amacrine types like Am1 with such an extended reach.

### Correspondences with molecular-morphological types

#### Tm5

Tm5a, Tm5b, and Tm5c were originally defined by single-cell anatomy and Ort expression (Gao et al. 2008; Karuppudurai et al. 2014). Tm5a is cholinergic, the majority of the cells extend one dendrite into the distal medulla, and often has a “hook” at the end of its lobula axon. Tm5b is cholinergic, and many cells extend several dendrites into the distal medulla. Tm5c is glutamatergic and extends its dendrites up to the surface of the distal medulla. Three of our types are consistent with these morphological descriptions (Fig. 7a), and receive direct input from inner photoreceptors R7 or R8.

#### Dm8

Molecular studies previously divided Dm8 cells into two types (yDm8 and pDm8), depending on whether or not they express DIP*γ* (Menon et al. 2019; Courgeon and Desplan 2019). The main dendrites of yDm8 and pDm8 were found to connect with R7 in yellow and pale columns, respectively. Based on its strong coupling with Tm5a, our Dm8a likely has some correspondence with yDm8, which is also selectively connected with Tm5a (Menon et al. 2019; Courgeon and Desplan 2019). But it is not yet clear whether there is a true one-to-one correspondence of yDm8 and pDm8 with Dm8a and Dm8b. It is the case that Dm8a and Dm8b strongly prefer to synapse onto Tm5a and Tm5b, respectively.

However, Tm5a and Tm5b are not in one-to-one correspondence with yellow and pale columns. Rather, the main dendritic branch of Tm5a is specific to yellow columns, while the main dendritic branches of Tm5b are found in both yellow and pale columns (Karuppudurai et al. 2014).

The relation of Dm8a and Dm8b to yellow and pale columns will be analyzed when more accurate photoreceptor synapses become available. At the present time, we can already reexamine some previous assumptions. The molecular studies assumed that (1) Dm8 cells should be in one-to-one correspondence with columns, and inferred that the (2) the ratio of yDm8 to pDm8 should be 7:3, the same as the ratio of yellow to pale columns. The actual numerical evidence provided for these assumptions was not altogether conclusive (Menon et al. 2019; Courgeon and Desplan 2019).

In v783, there are fewer than 400 Dm8 cells. Even after adjusting for underrecovery, we can be confident that the true number of Dm8 cells is much less than the number of columns (800). So assumption #1 can be discarded. It follows that there is no reason to believe assumption #2 either.

### Other information about types

We have already discussed quite a few types with partial coverage. There are many other cases in which partial coverage can be related to connectivity with a boundary type that only exists in a particular region of the visual field.

The Sm family contains the most types with partial coverage. This along with very similar stratification profiles makes Sm the most challenging family for cell typing, and connectivity analysis is indispensable.

While most singleton types (e.g. Sm⇐TmY17, Sm⇒aMe5⇐aMe17b, Sm⇒TmY16, and Sm⇒MeTu1⇒Tm35) are full-field, Sm⇐L3⇒Tm5_d is a singleton covering the ventral field only. Sm⇐aMe4⇒Mi15 is a singleton with a dorsal dendritic arbor in M7 and full-field axonal arbor in M1.

Of the types with partial coverage that contain multiple cells, Sm⇐CB0566⇐MeTu2, Sm⇐MC65⇒TmY5a, Sm⇐MeTu3⇒MeTu2, Sm⇐MeTu4⇐MeTu1, Sm⇐MeTu4⇒CB0566, Sm⇐Mi15⇒MLt5, Sm⇐Mi9⇒Tm32, and Sm⇒TmY3 are dorsal-only. Ventral-only types include Sm⇐Mi9⇒CB0165, Sm⇐TmY5a⇒Tm7, and Sm⇐Y12⇒TmY10. Notably, Sm⇐Mi9⇒Tm32 resembles similar stratification to Mi3 in upper and lower boundaries of the M7 layer. However, Sm⇐Mi9⇒Tm32 exhibits an additional stratification within the M4 layer.

Sm⇐Tm31⇒Tm25 and Sm⇐MC66⇐Mi10 are types covering the ventral two thirds of the medulla. Sm⇐CB0081⇐PS123 leaves the dorsal and ventral thirds of the visual field uncovered.

Sm⇐IB029⇐MeTu1 is a trio of H-shaped polarized cells with dendrites at the anterior rim, and axons at the posterior rim. Sm⇐CB0566⇐MeTu2 consists of three sickle-shaped cells at the dorsal rim. Sm⇐IB029⇒Tm34 consists of nine polarized cells with sickle-shaped axons arching in the anterodorsal medulla, and dendrites branching sparsely in the anteroventral region.

Most Pm types consist of 10 to 100 cells. One exception is Pm14 (Pm⇒C2), a jigsaw pair of full-field cells stratifying in M8. Another exception is Pm12 (Pm⇐TmY5a), a pair of full-field cells stratifying in M9, and sending tendrils up into M8.

Li2 is a set of eight cells, Li12 is a jigsaw pair, and Li16 is a pair of full-field cells.

In addition to mALC1 and mALC2, there are two new full-field singletons Li⇒Y3 and Li⇒LC13⇒TmY17. The four cells in Li⇐mALC4⇒LT33 are almost full-field, but their shapes are irregular and interesting.

For example, one covers dorsal and ventral regions but not the equator.

Li18 has been split into three types. Li⇐TmY17⇒TmY10 covers the whole visual field, while the other two Li18 types have only partial coverage of the visual field. Li⇐Tm8b⇒aMe20 covers a dorsal region except for the dorsal rim. It is tangentially polarized, with the axon more dorsal than the dendrites. Both axon and dendrite point in the posterior direction, perpendicular to the direction of polarization. The dendrites are more thickly stratified than the axon. Li⇐mALC5⇒Tm16 has ventral coverage only. The axons are in one layer, and extend over a larger area than the dendrites, which hook around into another layer and are mostly near the ventral rim.

LPi14 (LPi⇐T5a⇒H2) was called LPi1-2 by (Shinomiya et al. 2022), and is a jigsaw pair of full-field cells (Fig. 5f). LPi02 (LPi⇐T5a⇒LPLC2) is a more numerous type of smaller cell with similar stratification (Fig. 5g).

Four LPi types receive input from T4b and T5b. Two of them, LPi⇐T5b⇒T5b and LPi⇐T5b⇒TmY15, are reciprocally connected with T4b and T5b. LPi⇐T5b⇒T5b is a singleton full-field cell, which likely corresponds with LPi2b of (Shinomiya et al. 2022). LPi⇐T5b⇒CH is another singleton with output onto LPTCs that prefer front-to-back motion (CH and HS), as well as columnar cells with the same preference (T4a and T5a). LPi⇐T5b⇒CH. LPi⇐T5b⇒TmY17, the most numerous type in this group, has little or no output onto T4 and T5. It was called LPi2-1 by (Shinomiya et al. 2022).

Four LPi types receive input from T4c and T5c. The less numerous types LPi⇐T5c⇒LPLC4 and LPi⇐T5c⇒PLP127 each cover only part of the visual field. The strongest outputs of the ventral-only LPi⇐T5c⇒PLP127 are CB0203 and PLP127, VPNs with dendrites only in the ventral portion of the lobula plate. LPi⇐T5c⇒LPLC4 lacks coverage in the medial portion of the lobula plate. This type targets a subset of VS cells, the locations of which are related to the partial coverage. Both LPi⇐T5c⇒LPLC4 (Fig. 5h) and LPi⇐T5c⇒PLP127 are polarized in the tangential plane, showing a clear segregation between dendritic and axonal arbor. This is unlike most LPi types, which are amacrine. LPi⇐T5c⇒Y11 targets a number of columnar types, including columnar VPNs. The most numerous type in this group is LPi⇐T5c⇒VS, which was called LPi3-4 by (Shinomiya et al. 2022).

Three LPi types receive input from T4d and T5d. LPi⇐T5d⇒TmY14 is a full-field singleton that is reciprocally connected to T4d and T5d. LPi⇐T5d⇒LLPC2 and LPi⇐T5d⇒LPLC4 synapse onto columnar cells that target the central brain.

A final cell type is LPi⇐T5c⇐T5d, which receives roughly equal T4 and T5 input from the c and d preferred directions. This breaks the rule of input from one preferred direction, obeyed by all other LPi types. It is reciprocally connected to the same T4 and T5 types.

The full-field singletons (LPi⇐T5b⇒T5b, LPi⇐T5b⇒VCH, and LPi⇐T5d⇒TmY14) and the jigsaw pair (LPi⇐T5a⇒H2) are all predicted to be GABAergic. The remaining types contain tens of cells, and are mostly predicted to be glutamatergic.

